# CRISPR Screens Identify Novel Regulators of cFLIP Dependency and Ligand-Independent, TRAIL-R1-Mediated Cell Death

**DOI:** 10.1101/2022.08.17.504167

**Authors:** Neil Kuehnle, Scout Mask Osborne, Ziyan Liang, Mark Manzano, Eva Gottwein

**Affiliations:** Department of Microbiology-Immunology, Northwestern University, Feinberg School of Medicine, Tarry 6-735, Chicago, Illinois, 60611, USA; Mark Manzano, Department of Microbiology and Immunology, University of Arkansas for Medical Sciences, Little Rock, Arkansas, USA

## Abstract

Kaposi’s sarcoma-associated herpesvirus (KSHV) causes primary effusion lymphoma (PEL). PEL cell lines require expression of the cellular FLICE inhibitory protein (cFLIP) for survival, although KSHV encodes a viral homolog of this protein (vFLIP). Cellular and viral FLIP proteins have several functions, including, most importantly, the inhibition of pro-apoptotic caspase 8 and modulation of NF-κB signaling. To investigate the essential role of cFLIP and its potential redundancy with vFLIP in PEL cells, we first performed rescue experiments with human or viral FLIP proteins known to affect FLIP target pathways differently. The long and short isoforms of cFLIP and molluscum contagiosum virus MC159L, which are all strong caspase 8 inhibitors, efficiently rescued the loss of endogenous cFLIP activity in PEL cells. KSHV vFLIP was unable to fully rescue the loss of endogenous cFLIP and is therefore functionally distinct. Next, we employed genome-wide CRISPR/Cas9 synthetic rescue screens to identify loss of function perturbations that can compensate for cFLIP knockout. Results from these screens and our validation experiments implicate the canonical cFLIP target caspase 8 and TRAIL receptor 1 (TRAIL-R1 or TNFRSF10A) in promoting constitutive death signaling in PEL cells. However, this process was independent of TRAIL receptor 2 or TRAIL, the latter of which is not detectable in PEL cell cultures. The requirement for cFLIP is also overcome by inactivation of the ER/Golgi resident chondroitin sulfate proteoglycan synthesis and UFMylation pathways, Jagunal homolog 1 (JAGN1) or CXCR4. UFMylation and JAGN1, but not chondroitin sulfate proteoglycan synthesis or CXCR4, contribute to TRAIL-R1 expression. In sum, our work shows that cFLIP is required in PEL cells to inhibit ligand-independent TRAIL-R1 cell death signaling downstream of a complex set of ER/Golgi-associated processes that have not previously been implicated in cFLIP or TRAIL-R1 function.

## Introduction

Primary effusion lymphoma (PEL) is a non-Hodgkin lymphoma comprised of KSHV-transformed B cells that typically accumulate in body cavities^1, 2^. PEL is primarily seen in HIV/AIDS patients and has a poor prognosis, with an average survival below two years^3^. Although the etiology of PEL is incompletely understood, latently expressed KSHV oncoproteins are thought to play essential roles in the KSHV-mediated transformation of PEL cells, including the latency-associated nuclear antigen (LANA), viral interferon regulatory factor 3 (vIRF3), a KSHV-encoded D type cyclin (vCyc), and viral FLICE-inhibitory protein (vFLIP). Indeed, cultured patient derived PEL cell lines depend on the continued expression of several latency genes^4, 5^. vFLIP is considered a major candidate for a viral driver of KSHV-mediated B cell transformation in PEL due to its well-documented ability to activate NF-κB survival signaling^6^. A previous study by our lab explored the dependency of PEL cell lines on cellular genes using genome-wide CRISPR knockout screens^7^. This study identified a surprising dependency on cellular FLIP (cFLIP, encoded by *CFLAR*), despite the presence of the KSHV-encoded vFLIP locus. This study furthermore showed that only two of eight tested PEL cell lines depended on genes related to NF-κB signaling. These unexpected findings prompted us to further investigate the role of cFLIP in the context of KSHV-transformed PEL cell lines and to differentiate it from that of vFLIP.

Canonically, FLIP proteins act as dominant-negative inhibitors of caspase 8 (*CASP8*)-induced apoptosis by preventing formation of the death-inducing signaling complex (DISC) or directly inhibiting active CASP8^8–11^. Non-canonical cFLIP activities have been identified, including the regulation of NF-κB signaling^6, 12^. While KSHV vFLIP is a strong activator of NF-κB signaling, it is currently unclear whether vFLIP is a potent inhibitor of CASP8^13–15^. CASP8 is a pro-apoptotic protein in the cysteine protease (caspase) family that typically acts as the initiator caspase in the extrinsic apoptotic pathway induced by the death ligands tumor necrosis factor *α* (TNF*α*), Fas ligand, or TNF-related apoptosis-inducing ligand (TRAIL) (as reviewed in ^16^). The binding of these ligands to their specific death receptors initiates the formation of the death-inducing signaling complex (DISC), containing Fas-associated death domain (FADD) protein and inactive pro-CASP8. Autocatalytic cleavage of pro-CASP8 in DISC generates active CASP8, which initiates apoptosis. However, additional cellular processes converge on CASP8 to regulate cell death, including necroptosis, pyroptosis, autophagy-dependent cell death, and hyperactivation of the ER stress pathway/the unfolded protein response (UPR) ^17–22^.

Here, we sought to clarify the respective roles of cFLIP and vFLIP in PEL. Our results show that vFLIP is robustly expressed only in a subset of PEL cell lines, which also depend on genes that function in NF-κB signaling. The tight correlation of vFLIP expression with NF-κB dependency in PEL cell lines suggests that vFLIP functions in its well-established role as an activator of the NF-κB pathway in PEL. vFLIP cannot fully compensate for the absence of cFLIP, pointing to a non-redundant function of cFLIP in PEL cells. Using genetic screening and validation experiments, we established that cFLIP is required to inhibit ligand-independent, constitutive death signaling by one of the TRAIL receptors, TRAIL-R1 (encoded by *TNFRSF10A*).

## Results

### vFLIP expression is variable in PEL cell lines

We first examined the expression of human cFLIP and cyclin D2 and their viral homologs vFLIP and vCyc across 8 PEL cell lines and the KSHV-negative B cell line BJAB. As a control, we analyzed KSHV vIRF3, which is latently expressed in PEL from a distinct KSHV locus^4^. We observed comparable expression of the major isoforms of cFLIP, cyclin D2, and KSHV vIRF3 across the PEL cell lines (Fig. 1A). In contrast, vFLIP and vCyc expression varied greatly, with vFLIP expression detected only in BC-1 and BC-3. Importantly, these are the same two PEL cell lines shown to depend on NF-κB signaling genes in our previous study of cellular gene dependency in PEL cells (Fig. 1A, bottom) ^7^. vCyc expression mirrored the expression of vFLIP, as expected due to their bicistronic expression^23^, with the surprising exception of BC-2. Serial dilution of BC-3 lysates suggested that our Western protocol is sensitive enough to detect vFLIP and vCyc expression levels that are ∼20-50 fold below those of BC-3 (Fig. S1A, B). The observed patterns of vFLIP expression and NF-κB dependency are consistent with the published role of vFLIP in the activation of NF-κB survival signaling^6, 15, 24–27^, in the subset of PEL cell lines where vFLIP is expressed.

**Fig. 1.**
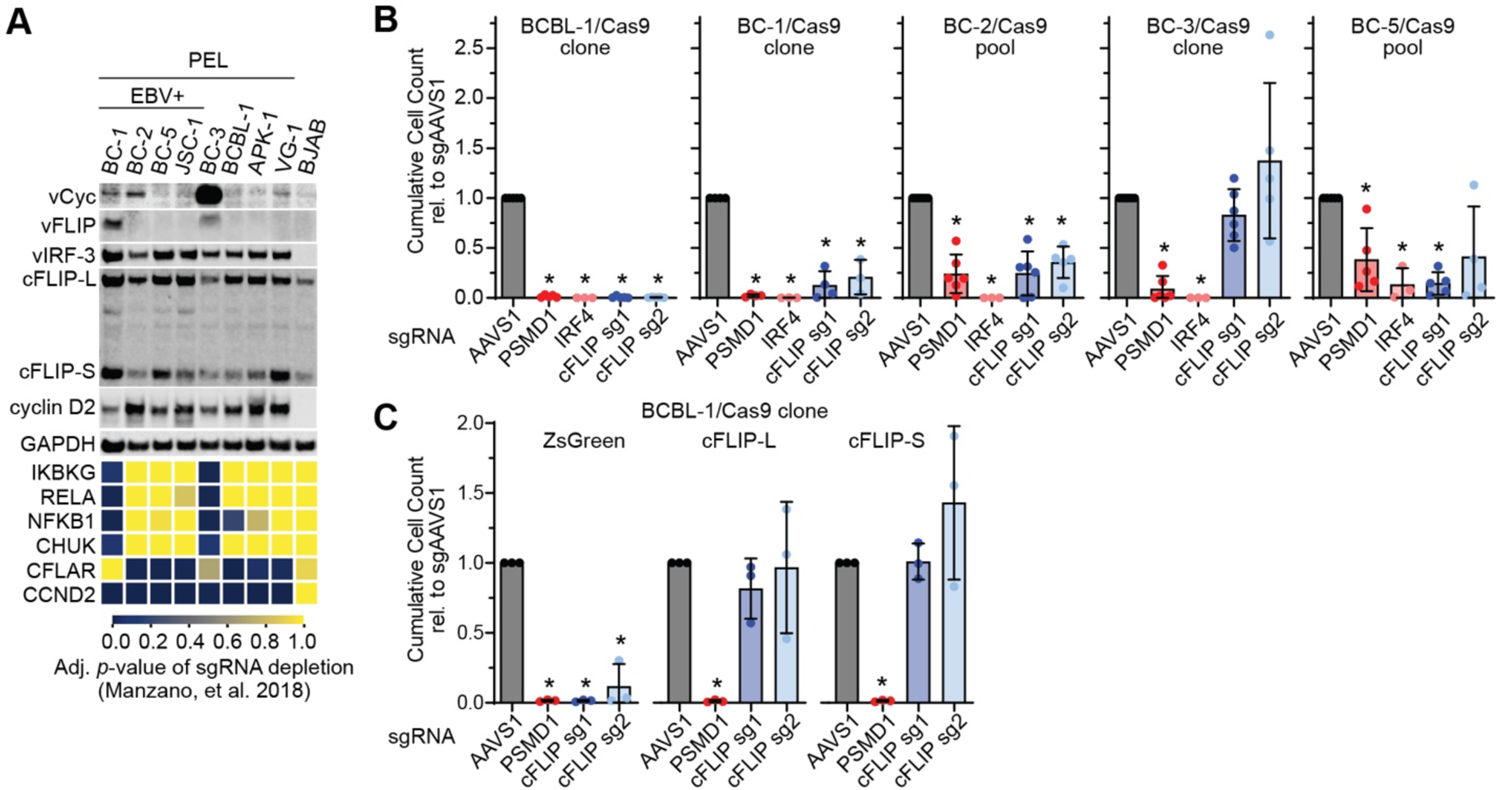
cFLIP is broadly essential in PEL cell lines. **A.** Top: Cell lysates from the indicated PEL cell lines or KSHV-negative BJAB cells were analyzed by Western blotting for the indicated proteins. The cFLIP splice variants cFLIP-L and cFLIP-S are marked. Bottom: Heatmap of depletion for sgRNAs targeting the indicated genes, using data from Manzano, et al. 2018. Lower FDR-adjusted *p*-values of depletion indicate significant sgRNA depletion in the screens, suggesting essential or fitness roles. **B.** Cas9-expressing PEL cell lines were transduced with the indicated sgRNAs at MOI 3 and selected with puromycin. Graphs show the endpoints (day 8-10) of cumulative cell growth curves relative to an sgRNA targeting the safe harbor locus AAVS1 (see Material and Methods). sgRNAs targeting PSMD1 and IRF4 are controls for other essential genes. Error bars represent SD (n = 4-6 independent repeats). For Western Blot control, see Fig. S1C. **C.** BCBL-1/Cas9 expressing ZsGreen, sgRNA-resistant cFLIP-L, or sgRNA-resistant cFLIP-S were challenged with the indicated sgRNAs. Error bars represent SD (n = 3 independent repeats). For Western Blot control, see Fig. S1E. In panels B and C, statistical significance for loss of cell viability compared to sgAAVS1 was analyzed using one-sided, one-sample t-testing (* denotes FDR-adjusted p <= 0.05), FDR-adjusted p-values are listed in Table 6. Rescue in panel C was significant as determined using a one-sided, independent two-sample t-test, with FDR-adjusted p-values listed in Table S6.

### cFLIP is broadly essential in PEL cell lines

We previously validated cFLIP dependency of the PEL cell line BCBL-1^7^. To establish cFLIP dependency more broadly across PEL cell lines, we performed single guide RNA (sgRNA)-induced functional inactivation (KO) of cFLIP followed by cumulative growth curve analysis relative to an sgRNA that targets a safe harbor locus (sgAAVS1) in BCBL-1/Cas9 and 4 additional Cas9-expressing PEL cell lines (Fig. 1B, Fig. S1C). sgRNAs against the proteasomal subunit PSMD1 and the transcription factor IRF4 served as positive controls for well-established essential genes in PEL cells. The cFLIP sgRNAs lack homology to vFLIP and did not affect vFLIP expression in BC-3 cells (Fig. S1D). For BC-1, which did not show significant depletion of cFLIP-specific sgRNAs in our original CRISPR knockout screens (Fig. 1A, ^7^), the experiment was done in a Cas9 cell clone with improved gene editing compared to the original cell pool^7^. Results showed that in addition to BCBL-1, BC-1, BC-2, and BC-5 require cFLIP for viability. Thus, BC-1 was false negative for cFLIP dependency in our original CRISPR screens^7^. In contrast, cFLIP single gene KO did not significantly affect the viability of BC-3 cells. Cell death following cFLIP KO in BCBL-1/Cas9 cells was rescued by lentiviral re-expression of either major cFLIP isoform (cFLIP-L and S; Figs. 1C, S1E). These data collectively confirm that cFLIP is essential in the majority of PEL cell lines. The essentiality of cFLIP in BC-1 furthermore suggests that expression of vFLIP may not be sufficient to overcome the requirement for cFLIP.

### cFLIP and vFLIP have distinct roles in PEL cell lines

To further investigate whether vFLIP and cFLIP are redundant, we tested if KSHV vFLIP can rescue cells from cell death after cFLIP KO. We additionally tested the viral FLIP proteins encoded by molluscum contagiosum virus (MCV), MC159L or MC160L, given that KSHV vFLIP and MCV MC159L and MC160L each have distinct functional activities (Fig. 2A). While either major isoform of cFLIP can inhibit CASP8 and activate NF-κB signaling in ectopic contexts^12, 28–30^, reports of KSHV vFLIP’s ability to strongly inhibit CASP8 are mixed or ambiguous. Conversely, MCV MC159L and MC160L both repress NF-κB signaling and only MC159L inhibits CASP8^8, 31–33^. Thus, this approach allows us to assess the redundancy of vFLIP with cFLIP and the importance of NF-κB activation and CASP8 inhibition for cFLIP dependency. Interestingly, we observed only partial rescue of cFLIP dependency after overexpression of KSHV vFLIP (Figs. 2B, C, S2A). In contrast, overexpression of MC159L conferred complete rescue, while overexpression of MC160L had no effect. Expression of each of the viral FLIP proteins was comparable to overexpressed cFLIP and did not affect the expression of endogenous cFLIP (Figs. S2A, B). KSHV vFLIP was furthermore overexpressed well above levels of the endogenous viral protein in BC-3 (Fig. S2C). Thus, the inability of vFLIP to efficiently rescue cFLIP expression is not due to lower expression relative to cFLIP or endogenous vFLIP. Collectively, these results suggest that FLIP proteins that are known potent inhibitors of CASP8 (cFLIP and MC159L), but not KSHV vFLIP, can efficiently rescue cFLIP dependency in PEL cells. Therefore, the essential role of cFLIP in PEL cells is substantially distinct from that of vFLIP.

**Fig. 2.**
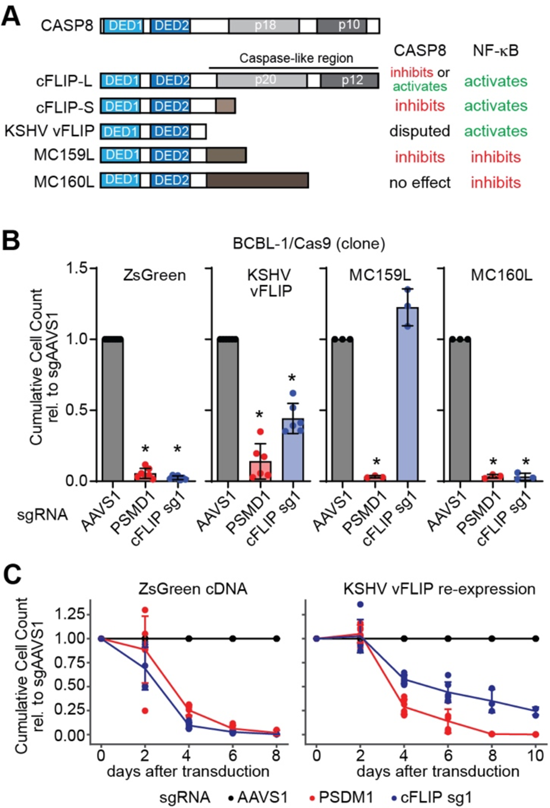
cFLIP and KSHV vFLIP are functionally distinct in PEL cell lines. **A.** Schematic representation of human CASP8, the cFLIP-L/S splice variants, KSHV vFLIP, and the MCV FLIP proteins MC159L and MC160L. Also shown are the characterized activities of each protein towards CASP8/the extrinsic apoptosis pathway and the NF-κB pathway. **B.** BCBL-1/Cas9 expressing ZsGreen, KSHV vFLIP, MCV MC159L, or MCV MC160L were challenged with the indicated sgRNAs. Analyses of cumulative growth curve experiments on day 6 after sgRNA transduction are shown. Error bars represent SD (n = 3 independent repeats). For Western Blot controls, see Fig. S2. Statistical significance for loss of cell viability compared to sgAAVS1 was analyzed using one-sided, one-sample t-testing (* denotes FDR-adjusted p <= 0.05). Rescue by expression of KSHV vFLIP and MCL159L was significant as determined using a one-sided, independent two-sample t-test. FDR-adjusted p-values are listed in Table S6. **C.** Extended growth curves for the ZsGreen and KSHV vFLIP data in panel B show that vFLIP rescue is significant, but not efficient. For details on statistical analysis see Table S6 and Material and Methods section.

### Genome-wide rescue screens uncover an ER/Golgi/CASP8-dependent death signaling program repressed by cFLIP

cFLIP has a well-characterized role in protecting cells from the effects of cell death ligands of the extrinsic apoptosis pathway^9, 11^. Indeed, Epstein-Barr virus (EBV)-immortalized lymphoblastoid cell lines (LCLs) depend on cFLIP expression for protection from autocrine TNF-*α* signaling^34^. TNF-*α* is not expressed in PEL cell lines (Fig. S3), however. We therefore sought to identify mechanisms underlying cFLIP dependency of PEL cell lines in genome-wide CRISPR/Cas9 rescue screens, outlined in Fig. 3A. In these screens, sgRNAs targeting genes whose inactivation overcomes cFLIP dependency should be enriched after toxic cFLIP inactivation and may represent putative components of the cell death process that is inhibited by cFLIP in PEL cells. We initially chose the background of BCBL-1/Cas9, which has strong cFLIP dependency and excellent CRISPR/Cas9 editing efficiency. Briefly, we transduced BCBL-1/Cas9 with the genome-wide Brunello sgRNA library^35^ and passaged the cell population to allow dropout of sgRNAs targeting essential genes. We next challenged the resulting cell pool with another sgRNA vector targeting either cFLIP or the AAVS1 safe harbor locus and further passaged the culture until a cFLIP-sgRNA resistant population was obtained (Fig. S4A). In an independent experiment, we used three separate shRNAs for toxic cFLIP knock-down alongside two negative control shRNAs targeting Renilla luciferase (RLuc) (Fig. S4B). We finally repeated the sg-cFLIP resistance screen in BC-2, which has robust cFLIP dependency (Fig. 1A, B). In each case, resulting cell pools were subjected to next generation sequencing of the Brunello sgRNA inserts and sgRNA composition was analyzed using MAGeCK’s robust ranked aggregation (RRA) (Table S1). Independent shRNAs targeting either cFLIP or RLuc were treated as replicates in this analysis. Results from the BCBL-1/Cas9 screens were more robust than those from the BC-2 screen, which was noisy, most likely due to relatively poor editing in the BC-2 Cas9 cell pool (Fig. S4C). Within BCBL-1, top hits from the shRNA-based challenge were statistically more significant but had less dramatic sgRNA fold-changes (Fig. S4C). This is likely due to the inclusion of three different shRNAs targeting cFLIP, each resulting in less complete toxicity than the single highly efficient cFLIP sgRNA (Fig. S4A, B). To account for these differences, we initially compared the BCBL-1 sgRNA and shRNA screens using a rank-based filtering approach. Intersecting the top 150 hits by RRA score from the sgRNA and shRNA challenge in BCBL-1 yielded 23 high-confidence hits (Fig. 3B, Table S2).

**Fig. 3.**
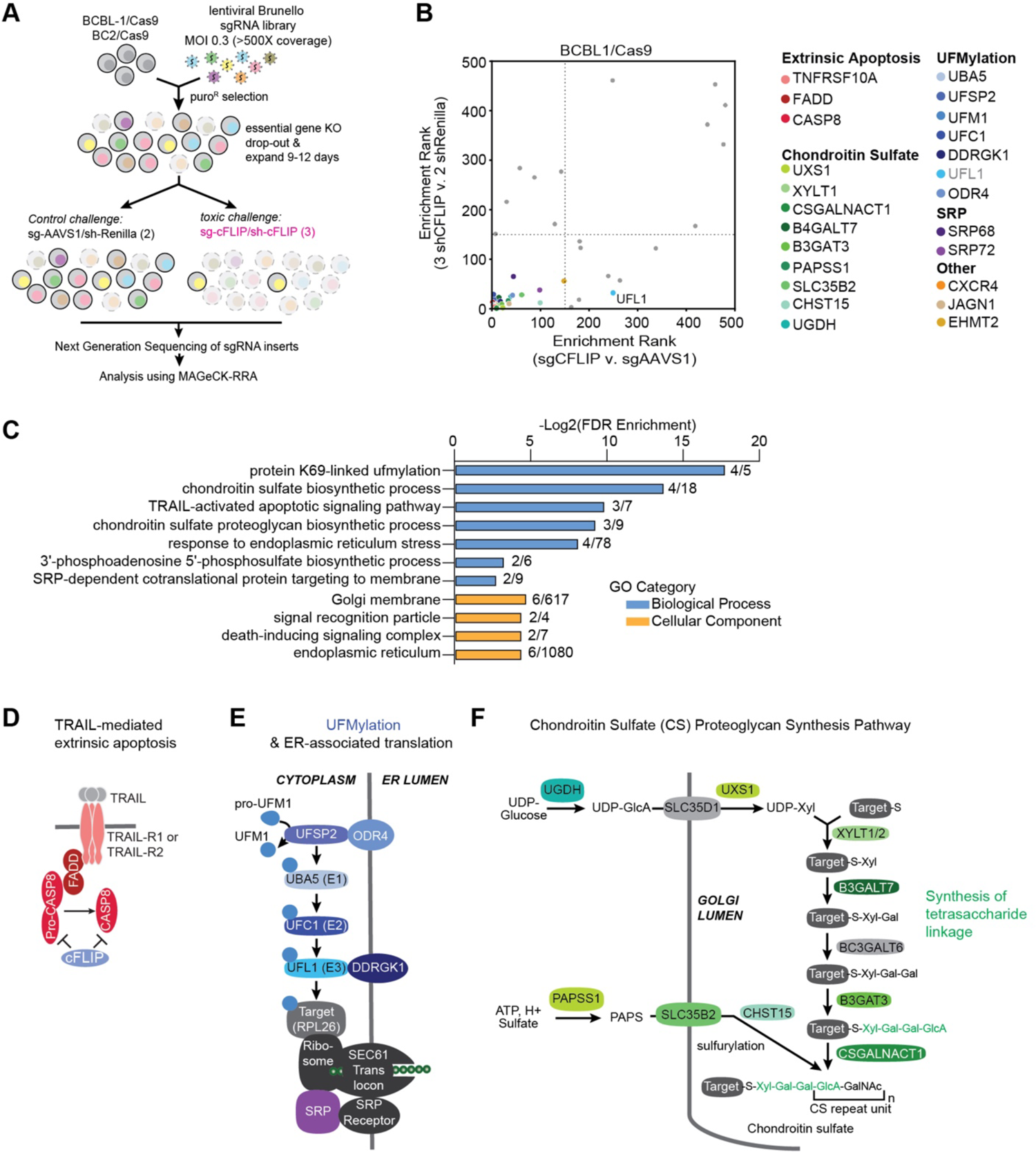
Genome-wide synthetic rescue screens uncover components of the death program repressed by cFLIP in PEL cells. **A.** Experimental design for CRISPR-based synthetic rescue screening. **B.** Results of the BCBL-1 resistance screens. Shown are the gene-level ranks output by MAGeCK-RRA based on library sgRNA enrichment after the toxic cFLIP shRNA challenge (y-axis) or the toxic cFLIP sgRNA-challenge (x-axis) relative to matched negative control perturbations. For clarity, only the top 500 hits in each screen are displayed. UFL1 was highlighted as a UFMylation pathway gene but does not pass our analysis cutoff. **C.** DAVID pathway analysis of the 23 high confidence hits obtained by intersecting the top 150 hits each from the BCBL-1 sgRNA and shRNA screens. Numbers at right show number of hits from each pathway. For complete DAVID output, see Table S3. **D.** Diagram of canonical death signaling via TRAIL through TRAIL receptors. Hits are colored as in panel B. **E.** Diagram of the UFMylation and ER-associated translation pathways, screening hits are colored as in panel B. UFM1 is cytoplasmic, but the UFM1 conjugation machinery is anchored on the cytoplasmic face of the ER membrane by adaptors ODR4 and DDRGK1 (UFBP1). RPL26 is the main characterized substrate for UFMylation and impacts translational pausing during ER-associated degradation. The SRP complex is involved in protein translocation and initiates translational pausing. **F.** Diagram of the Chondroitin sulfate (CS) proteoglycan synthesis pathway, screen hits are colored as in panel B.

DAVID pathway analysis^36^ shows that these high confidence hits function in three main pathways (Table S3, Fig. 3C): the extrinsic apoptosis pathway (including CASP8, FADD, and TNFRSF10A, encoding TRAIL receptor 1, Fig. 3D), protein modification by the ubiquitin-like modifier UFM1 (UFMylation, Fig. 3E), and chondroitin sulfate proteoglycan synthesis (Fig. 3F). Hits were furthermore enriched for genes associated with the ER and Golgi cellular compartments (Fig. 3C). Specifically, TRAIL-R1 and CXCR4 are transmembrane receptors which traffic through the ER-Golgi network. The UFMylation pathway mediates ubiquitin-fold modified 1 (UFM1) conjugation to lysine residues of target proteins at the cytoplasmic surface of the ER membrane^37–43^. SRP68 and SRP72 are components of the signal recognition peptide (SRP) complex, which cooperates with the SEC61A translocon to aid insertion of nascent transmembrane proteins into the ER^44–46^ (Fig. 3E). The SRP is involved in translational halting during membrane insertion^46, 47^. ER translocation of translationally paused nascent peptides requires UFMylation of the 60S ribosomal protein L26 (RPL26), which has been proposed as the primary target of UFM1 modification^48, 49^. Nine of the 23 high confidence hits participate in the chondroitin sulfate proteoglycan synthesis pathway (Fig. 3F), a process that occurs in the Golgi apparatus and involves the addition of chondroitin sulfate moieties to transmembrane proteins^50^. Jagunal homolog 1 (JAGN1) is required for vesicular transport from the ER or Golgi, although the exact step JAGN1 participates in is unclear^51^. Lastly, EHMT2 is a histone methyltransferase which has been implicated in the epigenetic regulation of the UPR and/or ER homeostasis^52, 53^. A subset of these genes also scored in the top 150 hits of the BC-2 screen, including CASP8 and TNFRSF10A (extrinsic apoptosis), UBA5 and UFSP2 (UFMylation), and UXS1, B4GALT7, CHST15, CSGALNACT1 (chondroitin sulfate proteoglycan synthesis), thereby confirming these genetic interactions of cFLIP in a second PEL cell line (Tables S1 and S2). Consistent with our screening results, BCBL-1 CASP8 KO pools derived using two different sgRNAs were completely protected from cell death after subsequent cFLIP sgRNA challenge (Figs. 4, Fig. S4D). These results confirm that our screens capture CASP8 as a key component of the cell death pathway that necessitates cFLIP expression in PEL cells.

**Fig. 4.**
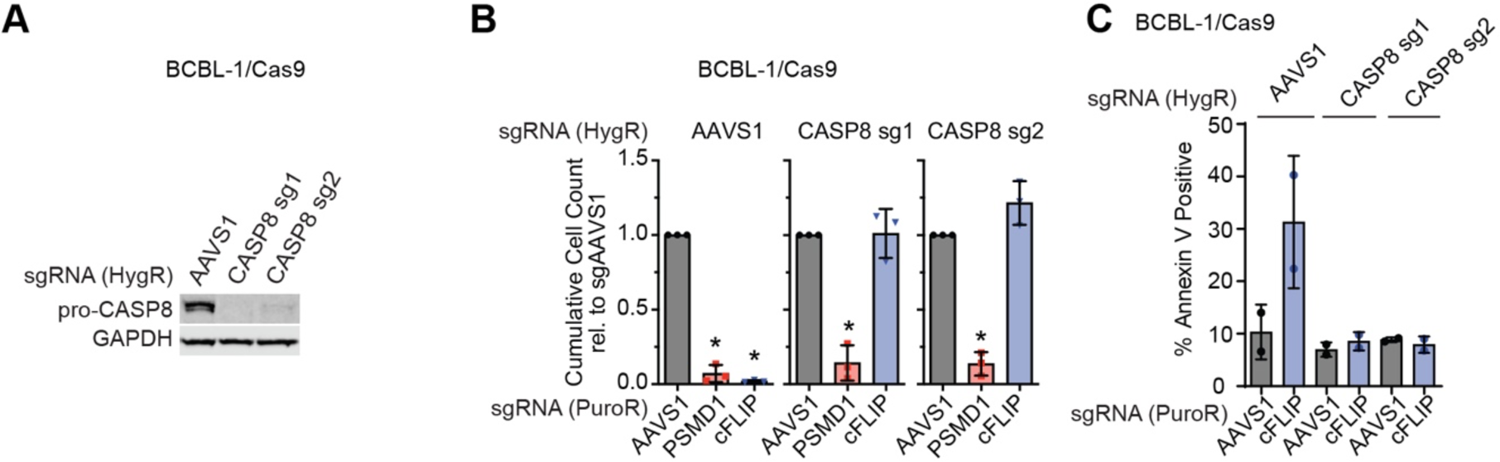
Cellular FLIP protects PEL cells from CASP8-induced cell death. **A.** BCBL-1/Cas9 cells were transduced with the indicated sgRNAs, selected with hygromycin, and knockout of CASP8 was confirmed by Western blot. **B.** The cell lines from panel A were challenged with puromycin resistant lentiviral sgRNA vectors at equal MOI and cumulative growth curve analyses were performed as previously. Error bars represent SD from 3 independent repeats. For Western Blot controls, see Fig. S4D. Statistical significance for loss of cell viability compared to sgAAVS1 was analyzed using one-sided, one-sample t-testing (* denotes FDR-adjusted p <= 0.05). Rescue by CASP8 KO was significant as determined using a one-sided, independent two-sample t-test. FDR-adjusted p-values are listed in Table S6. **C.** Cells were challenged with control or cFLIP sgRNAs as in panel B, but not selected with puromycin, and stained with Annexin V 48 hours after transduction (n=2 independent repeats, error bars indicate range).

### cFLIP protects PEL cells from ligand-independent TRAIL-R1-induced cell death

In addition to *CASP8*, *TNFRSF10A,* encoding TRAIL receptor 1 (TRAIL-R1/DR4) scored highly in our screens. Interestingly, neither the ligand for TRAIL-R1 (i.e. *TNFSF10* encoding TRAIL) nor *TNFRSF10B*, encoding TRAIL receptor 2 (TRAIL-R2/DR5), or other death receptors were hits in our screens (Table S1). As for CASP8, we generated TRAIL-R1 KO pools using two independent sgRNAs (Fig. 5A). Cumulative growth curve analyses following subsequent cFLIP KO confirmed that TRAIL-R1-deficient cells no longer require cFLIP expression for viability (Fig. 5B, Fig. S5A). TRAIL (ligand) mRNA is not well expressed in RNA-seq data from three different PEL cell lines, including BCBL-1 (Fig S3). Indeed, TRAIL ELISA on cellular supernatants or lysates of BCBL-1 and BC-3 cells failed to detect secreted or intracellular TRAIL^54, 55^ in either cell line (Fig. 5C). This assay detected levels of recombinant TRAIL well below the IC50 of “TRAIL-sensitive” cell lines within the literature^56^. BCBL-1 cells are furthermore TRAIL-resistant, even to hyper-physiological concentrations as high as 10 μg/mL (Fig. S5B). As additional controls, we generated TRAIL and TRAIL-R2 KO pools. While NGS-based sequencing of the targeted loci to characterize indel rates within our KO pools showed that >95% of each locus contains inactivating indels after editing (Table S4), genetic disruption of neither locus affected BCBL-1 dependency on cFLIP (Figs. 5B, S5A).

**Fig. 5.**
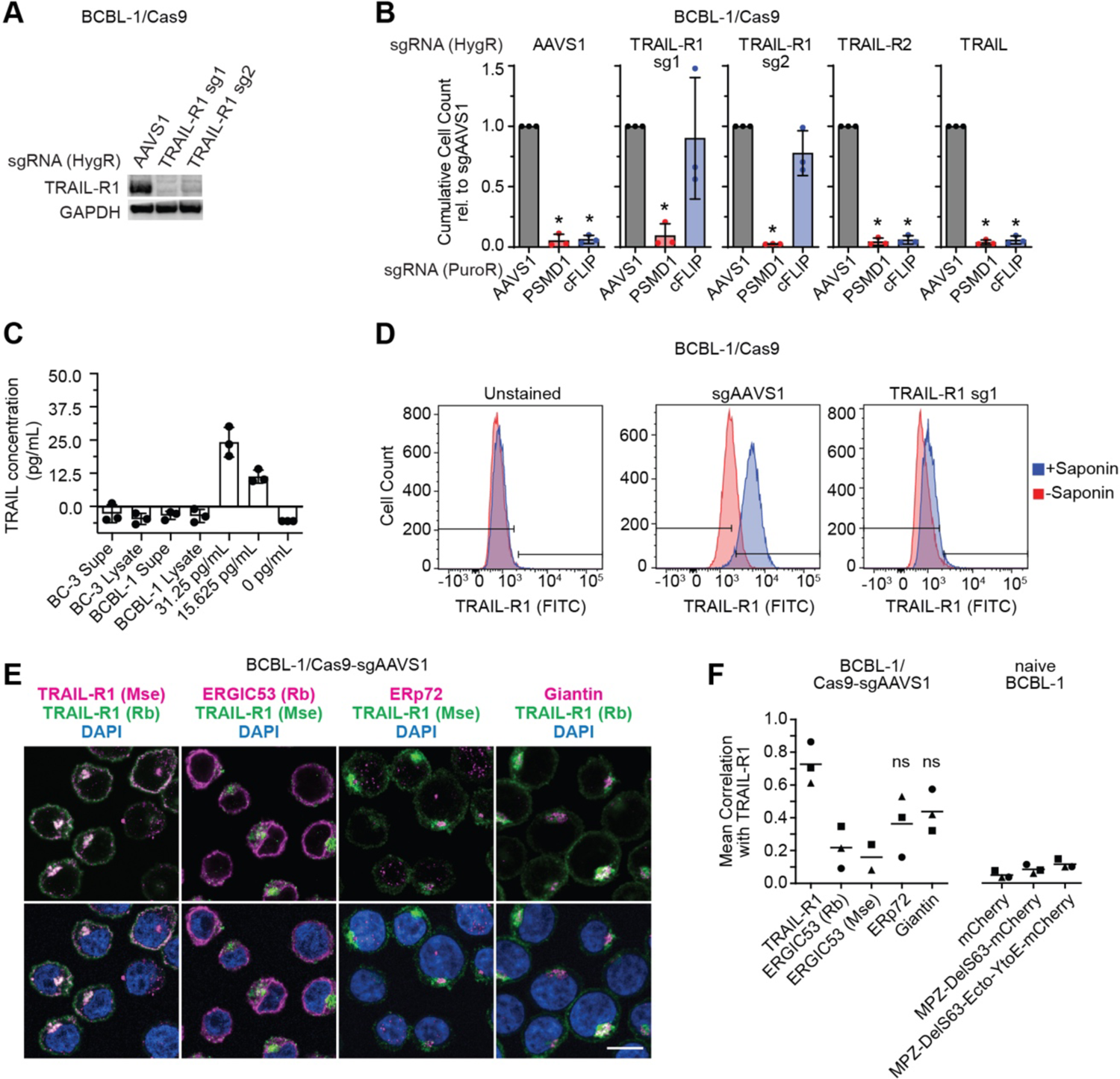
Cellular FLIP protects PEL cells from a ligand-independent, intracellular TRAIL-R1-mediated cell death program. **A.** BCBL-1/Cas9 cells were transduced with the indicated sgRNAs, selected with hygromycin, and knockout of TRAIL-R1 was confirmed by Western blot. **B.** Cell lines depicted in panel A were challenged with puromycin resistant lentiviral sgRNA vectors at equal MOI and cumulative growth curve analyses were performed as previously. Error bars represent SD from 3 independent repeats. For Western Blot control, see Fig. S5A. Statistical significance for loss of cell viability compared to sgAAVS1 was analyzed using one-sided, one-sample t-testing (* denotes p<= 0.05). Rescue by TRAIL-R1 KO, but not TRAIL-R2 or TRAIL KO, was significant as determined using one-sided, independent two-sample t-tests. FDR-adjusted p-values are listed in Table S6. **C.** BC-3 or BCBL-1 cells were plated at 2×10^5^ cells/mL, and supernatants or lysates were harvested 3 days later and used for anti-TRAIL ELISA assay. Known concentrations of recombinant human TRAIL confirm sensitivity down to the ∼10 pg/mL range. **D.** Knockout cell lines were fixed in 4% FA and stained for TRAIL-R1 with or without permeabilization by saponin. Shown at left are unstained control cells. Quantification over 3 independent repeats can be found in Fig. S6C. **E.** Representative images to establish colocalization of TRAIL-R1 with itself, using a positive control antibody directed against a different epitope, the ERGIC marker ERGIC53, the ER marker ERp72, and the Golgi marker Giantin. TRAIL-R1 is shown in green with other markers in magenta to depict overlapping signal as white. See Fig. S5E for results from a second ERGIC53 antibody and TRAIL-R1 KO cells. Scale bar = 10μm. Mse and Rb indicate antibodies raised in mice or rabbits, respectively. **F.** 3D colocalization of TRAIL-R1 with the indicated proteins was quantified using Pearson’s correlation coefficients (PCCs, n=3 independent repeats, except n=2 for mouse anti-ERGIC53). Representative images are shown in Figs. 5E, S5D-E. We quantified an average of ∼1800 cells per replicate. Statistical significance of the difference from TRAIL-R1 self-colocalization detected with two different TRAIL-R1-specific antibodies was determined by two-sided, independent two-sample t-test (* <=0.05). All markers other than ERp72 and Giantin (indicated by ns) were significantly less colocalized with TRAIL-R1 than TRAIL-R1 self-colocalization.

Ligand-independent activation of TRAIL-R2, and in some cases TRAIL-R1, has previously been described within the context of Golgi and ER stress-induced cell death^22, 57–61^. Ligand independent signaling of TRAIL receptor was proposed to follow the intracellular accumulation of TRAIL-R1 in the Golgi or TRAIL-R2 in the ER-Golgi intermediary compartment (ERGIC) ^61, 62^. Similarly, TRAIL-R1 in BCBL-1 is exclusively expressed intracellularly in PEL cells (Figs. 5D, S5C). TRAIL-R1 in BCBL-1/Cas9-sgAAVS1 colocalized with the ER marker ERp72 and the cis-Golgi marker Giantin, but much less extensively with the ERGIC marker ERGIC53 (Fig. 5E-F, S5E). A recent study in HCT116 cells has shown that a misfolded mutant of myelin protein zero (MPZ-DelS63) binds to TRAIL-R2 in the ERGIC and consequently triggers TRAIL-independent cell death, while MPZ-DelS63-EctoYtoE misfolds but does not interact with TRAIL-R2^62^. TRAIL-R1 did not colocalize with either form of MPZ in BCBL-1 after their doxycycline (DOX)-inducible expression with a C-terminal mCherry tag (Fig. 5F, S5D). Neither protein furthermore impaired the viability of BCBL-1 cells over the course of several days (not shown). Based on these findings, we conclude that cFLIP protects PEL cells from an intracellular, ligand-independent, TRAIL-R1-mediated cell death program originating within the Golgi or ER.

### UFMylation and JAGN1 promote TRAIL-R1 expression in PEL cells

To extend validation of the resistance screens beyond TRAIL-R1 and CASP8, we chose two hits from the UFMylation pathway (UFM1, DDRGK1), two genes from the chondroitin sulfate proteoglycan synthesis pathway (UGDH, CHST15), JAGN1, and CXCR4 for single gene KO by two independent sgRNAs each. Generation of an SRP68 KO cell line failed due to the essentiality of this gene in PEL cells, which was unsurprising since all six subunits of the full SRP complex are pan-essential based on data from the Cancer Dependency Map (Table S2). Regardless, we speculate that the screens correctly captured a biologically relevant connection to the SRP, due to the functional link between the SRP and UFMylation of RPL26. The essentiality of the SRP complex likely also explains why only 2 of 6 SPR complex members scored in our screen (Fig. 3B). We did not attempt generating an EHMT2 KO cell pool, since EHMT2 is also pan-essential and likely essential in PEL cells^7^. Western Blot analysis confirmed disruption of UFMylation in UFM1 KO pools (Fig. S6A). Interestingly, DDRGK1 KO did not reduce the most prominent UFM1-modified bands, suggesting that these represent upstream UFMylation pathway intermediates and/or other DDRGK1-independent UFMylation events. We furthermore confirmed robust editing in all cell pools by indel sequencing (Table S4). We obtained significant rescue from lethal cFLIP KO challenge for every hit, as predicted by our screens (Figs. 6A, Fig. S6B).

**Figure 6.**
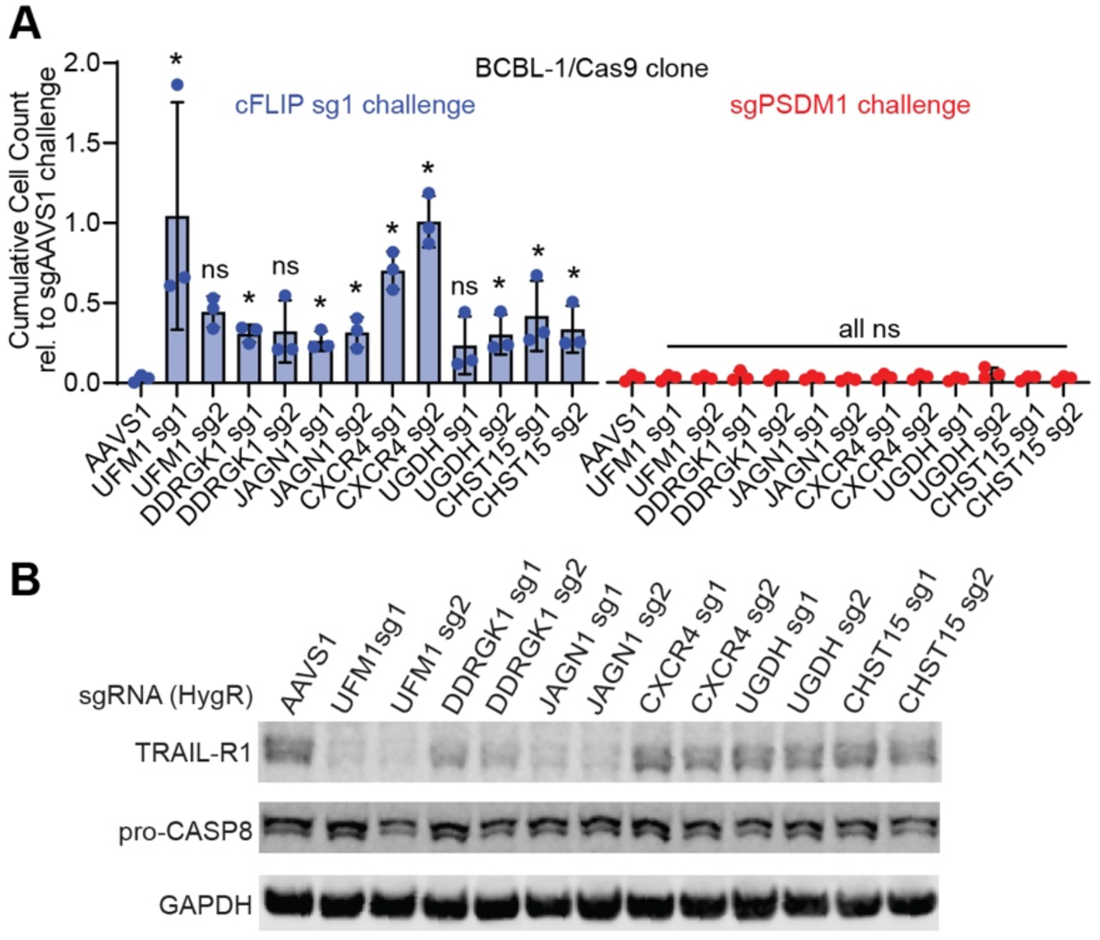
Knockdown of genes that participate in UFMylation, the chondroitin sulfate synthesis pathway, or of JAGN1 and CXCR4 overcome cFLIP dependency in PEL cells. **A.** Single-gene knockout pools were generated as previously for two genes each in the UFMylation and chondroitin sulfate biosynthesis pathway along with CXCR4 and JAGN1 and these cell lines were used to perform cumulative growth curves following transduction with a second sgRNA as previously. Statistical significance of rescue from cFLIP sg1 induced cell death was determined using one-sided, independent two-sample t-tests (* denotes FDR-adjusted p <= 0.05). FDR-adjusted p-values and statistical significance of loss of viability are listed in Table S6. Western controls are in Fig. S6B. **B.** Single-gene KO pools were analyzed for TRAIL-R1 and pro-caspase 8 expression by Western blotting.

UFMylation (UFM1 and DDRGK1) and JAGN1 were furthermore required for expression of TRAIL-R1, a type I transmembrane protein that contains an ER signal peptide (Fig. 6B). In contrast, KO of CXCR4 and chondroitin sulfate proteoglycan synthesis pathway genes (UGDH, CHST15) did not affect total TRAIL-R1 or CASP8 expression (Fig. 6B). Furthermore, none of the KOs we tested led to a redistribution of TRAIL-R1 to the cell surface from its typical intracellular location in PEL cells, in contrast to the recently reported redistribution of TNFRSF17 after SEC61A inhibition by others (Fig. S6C) ^63^. TRAIL-R1 mRNA levels in the knockout cell lines with reduced TRAIL-R1 protein levels were unchanged, indicating that reduced TRAIL-R1 expression is not due to changes in transcription or mRNA abundance (Fig. S6D).

While CXCR4 is best known for its role as a chemokine receptor and well expressed in PEL RNA-seq data, expression of its ligand, CXCL12, is not detectable in BC-1, BC-3, or BCBL-1 cells (Fig. S7A). To establish whether CXCR4 signaling is involved in the rescue phenotype, we tested if CXCR4 inhibitor (AMD3100) treatment overcomes the dependency on cFLIP, using BCBL-1 cells with Dox-inducible Cas9 expression^64^. In these experiments, CXCR4 inhibitor treatment failed to recapitulate the genetic CXCR4 KO effect (Fig S7B-C). This result and the lack of endogenous CXCL12 expression suggest that the role of CXCR4 in triggering TRAIL-R1-dependent cell death in PEL is unrelated to its normal signaling activity. In sum, our data suggest that UFMylation and JAGN1 are required for TRAIL-R1 expression, while CXCR4 and the chondroitin sulfate proteoglycan synthesis pathway may promote TRAIL-R1 signaling or affect cFLIP essentiality by other mechanisms.

## Discussion

Here we have distinguished the roles of KSHV vFLIP and human cFLIP in KSHV-transformed PEL cell lines. While cFLIP is required for its canonical function to inhibit CASP8 downstream of a death receptor signal, vFLIP was surprisingly not detected in all PEL cell lines and could not fully compensate for loss of cFLIP expression. Moreover, detectable vFLIP expression correlated with requirements for NF-κB-related genes, consistent with the well characterized role of KSHV vFLIP as an activator of NF-κB signaling. Thus, our data demonstrate that vFLIP and cFLIP have only limited functional overlap in a physiological context. Our results suggest that vFLIP has either low affinity to CASP8 or cannot effectively inhibit CASP8 once bound. Since we achieved very high levels of vFLIP overexpression in our rescue experiments, vFLIP sequestration from CASP8 by NEMO is less likely, although possible. These possibilities should be tested in future studies.

We identified TRAIL-R1, but not TRAIL-R2, as the critical death receptor that triggers CASP8-induced cell death in the absence of cFLIP. Unlike in EBV-transformed LCLs^34^, this process appears entirely ligand independent. Interestingly, such a ligand-independent process has been described in response to ER stress-induced cell death, mainly for TRAIL-R2. A ligand-independent role for TRAIL-R1 has also been reported, but primarily in the context of added Golgi and/or secretory stressors rather than general ER stress^22, 57–61^. Ligand independent cell death signaling by TRAIL receptors involves their intracellular accumulation and, similarly, TRAIL-R1 is detected within the ER and Golgi in PEL cells.

Additionally, our results were obtained without the use of ER/Golgi stress-inducing agents and instead represent a constitutively active TRAIL-R1 death signaling process in robustly growing cells. Importantly, PEL cells are of post-germinal center B cell origin and terminal B cell differentiation is typically accompanied by a broad expansion and alteration of the ER and Golgi. However, PEL cell lines typically lack B cell surface marker and immunoglobulin expression and the status of the ER/Golgi in PEL cell lines has not been investigated in depth. Interestingly, KSHV has been reported to activate the UPR and dampen its downstream transcriptional response during lytic reactivation and ER stress has been shown to trigger KSHV reactivation in latently infected cells^65, 66^. However, we did not detect colocalization of TRAIL-R1 with MPZ-DelS63 in latently infected BCBL-1, as has been reported for TRAIL-R2 in the context of cell death during the UPR in HCT116^62^.

In addition to TRAIL-R1, our work has implicated several other ER/Golgi resident processes in cFLIP dependency. High confidence hits in our cFLIP resistance screens included most genes of the UFMylation pathway. Inactivation of UFM1 or DDRGK1 reduced TRAIL-R1 expression, which is likely to explain their detection in our screens. Compared to ubiquitin, UFM1 modifies only few proteins. The 60S ribosomal subunit RPL26 has been proposed as the primary UFM1 target, where UFM1 modification promotes the processing of translationally stalled polypeptides at the ER^48, 49^. While we did not directly test the role of UFM1 or RPL26 in the translation of TRAIL-R1, UFMylation of RPL26 may explain the requirement for UFM1/DDRGK1 for TRAIL-R1 expression. UFMylation is induced in response to cellular stressors, directing nascent peptides to the ER for degradation^40, 48, 49, 67, 68^ and, accordingly, disruption of UFMylation has been reported to induce ER stress and increase cell death^67, 69, 70^. Of note, genes in the UFMylation pathway do not score as essential in our original PEL screens and their essentiality within the Cancer Dependency Map is mixed^7, 71^. Thus, it is plausible that UFMylation could similarly allow translation of pro-apoptotic proteins, like TRAIL-R1, to proceed. A varied requirement for UFMylation for cellular survival is further supported by findings that deletion of DDRGK1 in mouse plasma cells did not result in increased cell death^72^.

Like UFMylation, we observed that JAGN1 promotes expression of TRAIL-R1. While little is known about this gene, JAGN1 has been shown to participate in the secretory pathway in myeloid cells and antibody-producing B cells ^51, 73^. As with UFMylation, JAGN1 is increased in response to ER stress and its depletion has been linked to increased apoptosis in granulocytes^51, 74^. Importantly, our PEL CRISPR screens and establishment of a JAGN1 KO cell line suggest that JAGN1 is non-essential in PEL cells. Similarly, the Cancer Dependency Map indicates that JAGN1 is not broadly required for cellular survival^7, 71^.

Finally, our data suggest that chondroitin sulfate proteoglycan biosynthesis and CXCR4 are involved in the ligand-independent activity of TRAIL-R1. Since inactivation of neither pathway affected TRAIL-R1 expression or cause re-distribution of TRAIL-R1 to the cell surface, the exact mechanism underlying their link to TRAIL-R1 signaling and cFLIP dependency remains unclear. Interestingly, both CXCR4 and its primary ligand, CXCL12, have been identified as targets of chondroitin sulfate moieties, although the roles of these modifications remain unclear^75, 76^. However, CXCL12 is not endogenously expressed in PEL cells and TRAIL-R1-dependent cell death in PEL cells is not overcome by chemical CXCR4 inhibition. While there is a report of CXCR4-induced apoptosis, but this process was independent of CASP8 and could not be inhibited by MC159L, unlike the TRAIL-R1-dependent process that is inhibited by cFLIP in PEL cells^77^. Recent work indicates that misfolded integral membrane proteins, such as MPZ, can act as non-canonical ligands for TRAIL-R2 activity during ER stress-induced cell death^62^. Notably, rhodopsin, a member of the 7-pass transmembrane protein family to which CXCR4 belongs, also demonstrated this activity^62^. Future work should therefore test whether misfolding of endogenous CXCR4 serves as a trigger of ligand-independent TRAIL-R1 signaling in PEL cells. In sum, our data strongly support a role for TRAIL-R1 in incorporating multiple ER/Golgi-associated signals in PEL cells into a constitutive, ligand-independent cell death process that must be inhibited by cFLIP to allow for cellular viability.

## Materials and Methods

### Cloning

Sequences for oligonucleotides and synthetic DNA fragments can be found in Supplementary Tables S5A and S5B. All new vectors were confirmed by Sanger sequencing (ACGT, Inc.). pLenti SpBsmBI sgRNA Hygro (Addgene, 62205) constructs were cloned by ligating annealed oligonucleotides specified in Table S5C with the BsmBI-digested vector ^78^.

For rescue experiments, human C-terminally 3X-FLAG tagged FLIP long and short isoforms and KSHV vFLIP were ordered as codon-optimized dsDNA gene fragment blocks (Integrated DNA Technologies). This initial codon optimization introduced resistance to cFLIP sg1, while cFLIP sg2 resistance was introduced via two overlapping PCRs centered on the sg2 target site using the synthetic gene fragment as a template (primers 4672/4673 and 4674/4323). PCR products were then inserted between the AgeI and BamHI sites of vector pLC-IRES-HYG ^64^ using Gibson assembly. KSHV vFLIP was cloned from a codon-optimized gene fragment block containing the entire vCyclin/vFLIP locus using primers 4317/4318 and Gibson assembled with 3X-FLAG PCR products amplified from the cFLIP-S gene fragment (primers 4319/4323). PCR products containing MC159L (primers 4338/4339) and MC160L (primers 4340/4341) were amplified from pBabe-based vectors kindly provided by Dr. Joanna Shisler (UIUC) and Gibson assembled with 3X-FLAG PCR products amplified from the cFLIP-S gene fragment (primers 4342/4323 for MC159L or 4343/4323 for MC160L).

vCyc and vFLIP expression constructs for transfection into 293T in Fig. S1A were cloned into pMSCVpuro (Takara, Catalog # 631461) by PCR, using primers 3646/3647 (vCyc) and 3648/3649 (vFLIP), followed by Gibson assembly with the EcoRI/XhoI digested vector. LentiGuide-Puro expressing cFLIP sg2 was cloned by ligation of annealed oligos 4532/4533 with BsmBI-linearized vector as described ^7^.

The parental shR-miR expression vector pZIP-ZsGreen-T2A-Hyg-shNT4 was cloned by replacing the EcoRI-IRES-PuroR-NotI cassette of shERWOOD-UltramiR Lentiviral shRNA NT control #4 in pZIP-hCMV-ZsGreen-Puro (Transomic) with a PCR product containing a T2A-HygR cassette that was amplified using primers 3551/3552 and template lenti MS2-P65-HSF1_Hygro (Addgene Plasmid #61426). The initial control insert shRNA NT control #4 was replaced by digesting pZIP-ZsGreen-T2A-Hyg-shNT4 with NotI and MluI and inserting miR-30-based shRNAs for cFLIP (sh1, shM, shN) or negative control shRNAs targeting Renilla Luciferase (Ren.308 and Ren.713) using Gibson Assembly. These shRNAs were designed using SplashRNA and/or as previously published ^79^. dsDNA fragments specified in Table S5 were ordered and used for Gibson Assembly, either directly (shM, N) or after PCR-amplification with primers 4594/4607 (sh1, Ren.308, Ren.713).

To clone cFLIP sg1 for DOX-inducible gene inactivation (Fig. S7B, C), we replaced the sgAAVS1 sequence within pLX-sgAAVS1^64^ using overlap PCR. PCR fragment 1 was amplified using the forward primer 2430 and the sgRNA-specific reverse primer 5313. PCR fragment 2 was amplified using the sgRNA-specific forward primer 5312 and reverse primer 2431. Fragments 1 and 2 were joined using overlap PCR with primers 2430 and 2431, cut with XhoI and NheI and ligated with XhoI/NheI-digested pLX-sgRNA vector using T4 DNA ligase.

To clone the parental DOX-inducible lentiviral vector pCW-MCS-Zeo, we used Gibson assembly of a PCR fragment allowing for zeocin resistance (generated using primers 5327 and 5328) and SbfI/BamHI-digested pCW-MCS-PGK-P2A^80^. To clone pCW-Zeo-mCherry, pCW-Zeo-MPZ-S63Del-mCherry, and pCW-Zeo-MPZ-S63Del-EctoYtoE-mCherry, we performed Gibson Assembly of the AgeI-MluI vector fragment of pCW-MCS-Zeo with a PCR product containing mCherry (primers 5320/5321), gBlocks 5324 and 5322 (DelS63), or gBlocks 5326 and 5322 (DelS63/EctoYtoE).

### Western blotting

Cells were pelleted, washed in ice-cold PBS, in some cases stored at −80°C, and lysed for 30 minutes in ice-cold RIPA buffer containing 1X protease inhibitor cocktail (Sigma-Aldrich, catalog P8340). Lysates were then subjected to five cycles (30 seconds on/30 seconds off) of sonication at 4°C on high intensity in a Bioruptor Sonication System (Diagenode) and subsequently cleared by centrifugation at 14,000g for 15 minutes. Lysates were diluted 5- to 10-fold and quantified by Pierce BCA Protein Assay (ThermoFisher Scientific, 23225). Lysates were denatured by heating at 70°C for 10 minutes in 1X LDS buffer (ThermoFisher, NP0008) containing a final concentration of 2.5% beta-mercaptoethanol and loaded onto Bolt 4-12% Bis-Tris gels (ThermoFisher Scientific; NW0412) at equivalent concentrations (15-20 µg for cFLIP knockout validation based on available lysates; 50 µg for vCyc/vFLIP, and 20-30 µg for all other proteins). SDS-PAGE was performed in 1X MES buffer and proteins were transferred onto 0.2 µm nitrocellulose membranes (Bio-Rad, 1620112).

Membranes were blocked at room temperature for 1 hour and probed overnight at 4 °C with primary antibodies at the concentrations indicated in Table S5D. Images were captured on an Odyssey FC (LI-COR Biosciences) after incubation with IRDye 800CW secondary antibodies (LI-COR Biosciences) at a 1:15000 concentration. In some cases, anti-rat HRP (Santa Cruz) at a 1:5000 concentration was used for detection and developed using SuperSignal West Femto Maximum Sensitivity Substrate (ThermoFisher Scientific, 34094) following manufacturer protocol. Contrast/brightness was dynamically adjusted as needed using ImageStudio. Bands were quantified in ImageStudio Lite version 5.2 (LI-COR Biosciences). Original Western Data are included as one supplemental file.

### Cell culture

PEL and/or BJAB cell lines were maintained in RPMI 1640 with L-glutamine (Corning, MT10040CM) supplemented with 10% (BC-1, BCBL-1) or 20% (BC-2, BC-3, BC-5, BJAB) Serum Plus-II (Sigma-Aldrich, catalog number 14009C, lots 15H243 and 21C421), 10 µg/mL gentamycin (ThermoFisher Scientific, 15710064) and 0.05 mM beta-mercaptoethanol (Sigma-Aldrich, M3148-25ML). HEK293T cells were maintained in DMEM (Sigma-Aldrich, D5796) supplemented with 10% Serum Plus-II and 10 mg/mL gentamycin. PEL cells were counted by trypan exclusion and split every second day to approximately 2×10^5^ cells/mL (or 3×10^5^ for BC-5). 293T cells were maintained at visually sub-confluent densities and split every 2 days. PEL cell lines expressing lentiCas9-Blast (Addgene, 52962) were previously generated and validated by STR profiling^7^. The BC-1/Cas9 clonal cell line utilized in Fig. 1B was subcloned and selected for editing efficiency compared to the parental Cas9 BC-1 pool utilized in the screens. This BC-1/Cas9 clone was re-confirmed by STR profiling. A stable BCBL-1 cell clone allowing for DOX-inducible Cas9 expression (BCBL-1/pCW-Cas9) was reported previously^64^. For cellular growth curves and functional titration of lentivirus other than pZIP-based vectors (see below), cells were counted by flow cytometry using spike-in of a known amount of AccuCount Blank Particles 5-5.9 µm (Spherotech, ACBP-50-10) as previously described^7^.

### Cumulative cellular growth curves

Importantly, all lentiviral vectors used for growth curves analyses were titrated in naïve parental cell lines to ensure comparable titers between control sgRNAs and toxic sgRNAs. For growth curves, cells were plated at a density of 3×10^5^ cells/mL and transduced at the multiplicities of infection (MOIs) indicated in the text or figure legends. Medium was exchanged one day after transduction and cells were selected with 1.2 µg/mL puromycin for 2 days; 200 µg/mL hygromycin for 3 days (titrations) or 5-7 days (stable cell line production). Every second day, cells were counted by flow cytometry as described above and then split to a density of 2×10^5^ cells/mL. This process was repeated until cells that had received toxic sgRNAs were too sparse to split or for up to 10 days. Cumulative cell counts were calculated by taking the measured cell concentration at a given timepoint multiplied by the product of all previous dilution factors and normalized to cumulative cell counts obtained for the sgAAVS1 control.

Statistical testing for reduced cellular viability was performed using one-sided, one-sample t-testing (H0: P(µ=1) with HA: P(µ<1)). For rescue growth curve experiments (i.e. in the context of ectopic FLIP expression or a knockout cell pool) an additional one-sided, independent two-sample t-test was performed, comparing the relative cumulative cell counts for each guide between the experimental cell line and the control line (i.e. sgAAVS1-Hygro or ZsGreen-Hygro). In the case of ZsGreen or KSHV vFLIP expression for rescue experiments, where n= 3-6 were obtained for up to day 10 (Fig. 2C), ordinary least squares regression lines were fit. Cumulative cell counts relative to cell-line matched sgAAVS1 were used as the singular dependent variable with one or two predictor variables. To measure depletion, data was subset on the indicated cell line (ZsGreen or vFLIP expression) and sgRNA (sgPSM1 or cFLIP sg1) with the timepoint (>0 days after transduction) as a singular predictor variable. To measure rescue, data was subset on a particular sgRNA (sgPSMD1 or cFLIP sg1) with the cell line (ZsGreen or vFLIP) as an additional categorical predictor variable besides the timepoint (2-8 days post-transduction). A one-sided t-test was then used to determine whether the coefficients for day (depletion) or cell line (rescue) were < or > 0, respectively.

### Lentiviral production, infection, and titration

Except for pZIP-based vectors, lentivirus was produced using packaging plasmids psPAX2, pMD2.G, and transfer vectors at 1.3:0.72:1.64 molar ratios. HEK293T Cells were seeded at a density of 1.25×10^7^ per 15-cm dish and transfected with 30 µg of DNA using a 1:3 to 1:3.5 ratio of µg of DNA to µL PEI. Media were replaced after ∼6 hours and cell supernatants containing viral vectors were collected 48-72 hours later and filtered through 450nm pore size filters. Viral vector preparations were used as is or concentrated by ultracentrifugation or using Amicon Ultra Centrifugal Filter Units (Millipore, UFC910024) following manufacturer protocol. In some cases, lentivirus was stored at −80°C.

Lentiviral vector titers were measured using functional titration of serially diluted viral stocks. The multiplicity of infection (MOI) of each dilution point was estimated using the percentage of live cells relative to an untreated control and a resulting viral concentration (transducing units/mL) was calculated from the average of multiple dilution points assuming a Poisson distribution 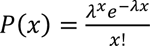, to estimate the mean event rate λ (MOI) assuming P(x=0) is the percentage of uninfected cells.

Transductions were carried out at 3×10^5^ cells/mL and the indicated MOIs under normal culture conditions supplemented with 5 µg/mL polybrene (Sigma-Aldrich, TR-1003-G). Culture medium was replaced ∼24 hours after transduction and antibiotic resistance selection was carried out using 1.2 µg/mL puromycin for 2 days; 200 µg/mL hygromycin for 3 days (titrations) or 5-7 days (stable cell line production and CRISPR screens). Procedures for pZIP-ZsGreen-T2A-Hygro shRNA vectors are described below.

### CRISPR-based single-gene knockout

CRISPR knockout was achieved by transduction of published Cas9-expressing cell pools or clones and published lentiviral single guide RNA constructs for AAVS1, PSMD1 sg1, IRF4 sg1, cFLIP sg1. A new cFLIP sg2 was cloned for this paper since sg2 from the previous study performed sub-optimally. The new cFLIP sg2 corresponds to sgRNA 2 (by rule set 2 score, on target activity) from the Brunello library. sgRNAs in pLentiGuide-Puro (Addgene, 52963) were used for cumulative growth curve knockout, while long-term single-gene knockout cell lines were generated using sgRNAs in pLenti SpBsmBI sgRNA Hygro (Addgene, 62205) containing a hygromycin resistance cassette. Sequences of all sgRNAs can be found in Table S5C.

### CRISPR-based synthetic rescue screens with sgRNA challenge

Library virus was generated and titrated as described above using the Brunello sgRNA pool library in lentiGuide-Puro (Addgene, 73178). In all cases, approximately 9.2×10^7^ to 1.5×10^8^ cells were transduced at an MOI of 0.3 to result in a single sgRNA per cell at a coverage of 300-500X cells per guide. Transduced cells were selected using 1.2 μg/mL puromycin, and dropout was allowed to proceed for 12 days with at least 3.6×10^7^ cells maintained to preserve library complexity.

The resulting cell pool was then challenged with either cFLIP sg1 or the negative control sgAAVS1 in pLenti SpBsmBI sgRNA Hygro at an MOI < 0.5. Cells were selected with hygromycin and cultured until resistance to cFLIP knockout was apparent by cellular proliferation, i.e. until day 12 after sgRNA transduction. At this point, approximately 3-7×10^8^ cells (500-1000X coverage) were harvested, washed in ice-cold PBS, and snap frozen.

### shRNA-based synthetic rescue screens

pZIP transfer vectors were packaged in Lenti-X 293T cells (Takara) using molar ratios of 0.2 transfer vector to 0.51 psPax2 to 0.29 pMD2.G. Briefly, 2.1×10^7^ pLenti-X 293T cells/15 cm dish were transfected with ∼48.5 μg total plasmid and supernatant was harvested 48 hours later without replacing media. Supernatants were filtered through 450nm pore size filters and concentrated by ultracentrifugation (Beckman SW32Ti, 25,000 rpm, 90 min), resuspend in OptiMEM on a platform shaker for at least 60 min at 4°C and followed by pipetting up and down 20 times, aliquoted, and snap frozen at −80°C. Virus aliquots were thawed at 37°C and cleared by centrifugation at 400 g for 5 min before use. Titration of the negative control vectors was performed based on ZsGreen expression and flow cytometry in BCBL-1/Cas9 clone C2. Titration of the toxic cFLIP shRNAs was performed in 293T cells lacking Dicer (293T/NoDice) ^81^, relative to a negative control with known titer in BCBL-1/Cas9. BCBL-1/Cas9 Brunello library transduction was performed at MOI 0.3 in eleven 15 cm dishes at 5×10^5^ cells/mL and 50mL/dish, ∼920x theoretical sgRNA coverage. Cells were selected using 1.2 μg/mL puromycin from days 1 to 4 after transduction and expanded. On Day 9 after Brunello transduction, seven 15 cm dishes of cells (3×10^5^ cells/mL and 50 mL/dish) were challenged with each shRNA vector at MOI 0.5. The next day, cells were collected by centrifugation and resuspended in fresh medium containing 200 μg/mL Hygromycin. Cell pools were maintained at coverage and optimal cell density with hygromycin for one week and at least 4×10^7^ cells/condition were harvested on day 12 into the shRNA challenge.

### CRISPR library preparation and next-generation sequencing

Genomic DNA was isolated following a previously described protocol and library preparation was performed as previously described for sgRNA and shRNA screens in BCBL-1 ^7^. Resulting libraries were quantified by Qubit (ThermoFisher Scientific, Q33231) prior to further QC and quantification by Bioanalyzer (Agilent) and Kapa qPCR (Roche). Multiplexing of sgRNA challenge screens in BCBL-1 was performed utilizing 6-bp indexes and sequenced on a single lane of a HiSeq4000 (Illumina) using 50bp single-end (SE) reads with 10% PhiX spike-in as previously described^7^. For sgRNA challenge screens in BC-2, sequencing was performed on a single flow-cell of an NextSeq 500 (Illumina) using 75bp SE reads with 10% PhiX spike-in due to the discontinuation of the HiSeq4000 platform. For these runs, library preparation was modified slightly to combine genomic amplification and addition of adapters/barcode into a single PCR step with custom 8bp indexes (sequences can be found in Table S5E). The shRNA-based screens were prepped and sequenced twice as a quality control using both approaches/platforms and reads were subsequently merged for further analysis. Demultiplexed reads have been deposited on GEO (accession GSE210445). 5’ adapters were trimmed from raw CRISPR sequencing reads and trimmed to length (20 bp) from the 3’ end using Cutadapt v4.1 ^82^. Reads were then aligned using Bowtie v1.1.2 9 ^83^. Guide-level counts were obtained using MAGeCK v0.5.9 and statistical testing was performed using MAGeCK’s robust rank aggregation (RRA) algorithm using median normalization ^84^.

### CRISPR Knockout Pools Indel Sequencing

A 200-300 bp region around each indicated guide was amplified from genomic DNA using the primers indicated in Table S5F and sent to Massachusetts General Hospital CCIB DNA Core for low depth sequencing. Raw FASTQ files were assembled into contigs by MGH and the resulting effect on the coding sequence was determined using custom alignment and classification script built using Biopython ^85^ to estimate indel, frameshift, and early stopping incidence. Briefly, assembled contigs were aligned to the coding sequences targeted by each amplicon to determine optimal positioning of the contig in reference to the full coding sequence. All alignments utilized a simple pairwise scoring matrix (+1 for matches, +0 for mismatches, −.99 for gap opening, and −0 for gap extensions) chosen by iterative manual assessment of the collective results for the most abundant contig for each gene. The resulting, fully assembled coding sequence was then classified as an indel or non-indel for all sequences. Indels were further characterized as in-frame or frameshifted based on the number of nucleotides lost or gained across all alignment gaps. Early stopping events for all variants were predicted by *in silico* translation of predicting coding sequences. Raw FASTQ output was also analyzed using CRISPResso2^86^ to obtain estimates of the number of modified and frameshifted reads.

### mRNA Sequencing

A frozen BC-1 lysate in Trizol was submitted to BGI Group (Shenzhen, China) for single-ended, polyA-enriched RNA sequencing. Un-trimmed reads were aligned to the Gencode 41 annotation of GRCh38.p13 with STAR v2.7.10. Raw sequencing data has been made available on GEO (accession GSE210446).

### RNA Purification and Reverse Transcription

5×10^6^ Cells were washed in ice-cold PBS, pelleted, and resuspended in 300 uL TRIzol (Invitrogen, 15596026). RNA was isolated using the Zymo Research Direct-zol RNA Miniprep Kit (Fisher Scientific, NC1057004) following the manufacturer’s protocol. Purified RNA was treated with RQ1 RNAse-free DNAse (Promega, M6101) using 1 μL DNase per 5 μL RNA in a final volume of 10 μL volume. Poly-dT primed reverse transcription was performed using the SuperScript First-Strand Synthesis System (Invitrogen, 11904018) using 1 μg of RNA as template in a 10 μL reaction, including water-only reactions as non-template controls, following manufacturer protocols including optional RNAse digestion after reverse transcription. Samples were then diluted to 100 ng calculated input RNA/4.5 μL prior to TaqMan assay.

### TaqMan Assay

Real-time PCR was performed by standard TaqMan Assay on either the QuantStudio 7 or Quantstudio 5 Real-Time PCR platform (Applied Biosystems). Briefly, a mastermix of 1 μL TaqMan probe (Applied Biosystems, Hs00269492_m1 or Hs00984230_m1) to 10 μL TaqMan Universal Master Mix II (Applied Biosystems, 4440043) was prepared for each probe. 5.5 μL of mastermix was added per well, followed by 4.5 μL (equivalent to 100 ng calculated input RNA) cDNA template. Three technical replicates were included, and 2^-ΔΔCT^ was calculated relative to the endogenous B2M control and sgAAVS1-transduced cell line for each biological replicate^87^. Biological replicates represent independently harvested cell pellets of the indicated cell lines. Statistical testing for differences between samples was performed using one-way analysis of variance. Post-hoc testing was performed using Dunnett’s method and sgAAVS1 as the control group.

### Inducible CRISPR/Cas9 Assay and AMD3100 Treatment

For data shown in Fig. 7B-C, we transduced BCBL-1 with stably integrated DOX-inducible Cas9 (BCBL-1/pCW-Cas9^64^) with pLX-sgRNA-lentiviral vectors expressing sgRNAs targeting AAVS1, PSMD1, or cFLIP sg1 at ∼MOI 2 and selected transduced cells with 7.5 μg/ml Blasticidin S (Sigma-Aldrich, SBR00022). We induced Cas 9 expression by treatment with 1 μg/ml Doxycycline hyclate (DOX, Sigma-Aldrich, D9891-1G) or left the cultures uninduced. At the same time, we further treated each cell line and DOX-treatment group with DMSO or 25 μM AMD3100 (Selleckchem, S8030). Cells were counted by trypan exclusion every 2-3 days, pelleted, and re-suspended to 2×10^5^ cells/mL in fresh medium containing the specified drugs.

### Cellular staining and flow cytometry

Cells were washed twice in ice-cold PBS and APC Annexin V (BD Biosciences, 550475) and 7-AAD (BD Biosciences, 559925) staining was performed on 1×10^5^ cells in 100 μL of 1X Annexin V binding buffer (BD Biosciences, 556454) following manufacturer recommended guidelines, using 5 μL of each stain for 15 minutes at room temperature in the dark. Necrotic cells were generated for Annexin V and 7-AAD compensation controls using a 30-minute heat shock at 95°C. Stained cells were diluted with 400 μL 1x Annexin V binding buffer and flow cytometry was performed on a BD FACSCanto II (BD Biosciences) within 1 hour.

For TRAIL-R1 staining, cells were harvested and washed twice in ice-cold PBS and resuspended in PBS at a concentration of 1×10^6^ cells/mL. 500 μL of cells were then fixed in PBS containing 4% w/v formaldehyde (FA, ThermoFisher Scientific, PI28908) for 10 minutes followed by two more PBS washes. On the final wash, cells were split into two and each resuspended in 250 μL PBS containing 1% w/v bovine serum albumin (BSA, Sigma, 10735078001) with or without the addition of 0.1% w/v saponin (Sigma-Aldrich, 47036). Cells were pre-incubated for 15 minutes at room temperature with 250 μL PBS/1% BSA with or without 0.1% w/v saponin, containing 1:400 of Human BD Fc Block (ThermoFisher Scientific, BDB564219) per condition, followed by addition of 1:100 (2.5 μL in 250 μL) of FITC-conjugated anti-TRAIL receptor 1 (Abcam, ab59047; specific to the extracellular domain of TRAIL-R1) for 30 minutes at room temperature in the dark. Cells were then washed 3X with PBS/1% BSA with or without 0.1% w/v saponin followed by a final re-suspension in 500 μL PBS containing 0.01% FA.

Stained cells were stored for up to 2 days at 4°C in the dark (TRAIL-R1) and flow cytometry was performed on a BD FACSCanto II (BD Biosciences). Raw data were exported to FlowJo v10.8.1 for analysis and plotting.

### BCBL-1 cell lines expressing MPZ-Del63-mCherry constructs

To allow for inducible expression of mCherry, MPZ-Del63-mCherry, or MPZ-DelS63-EctoYtoE-mCherry, we transduced naïve BCBL-1 cells at MOI <1. We selected transduced cells with 175 μg/ml zeocin (Life Technologies, R25005) and induced protein expression by treatment with 1 μg/ml DOX. Results in Figs. 5E-F and S5D-E are from day 3 after induction and three independent inductions.

### Immunofluorescence and colocalization analysis

Cells were collected by centrifugation, washed with PBS, and resuspended in PBS at 1×10^6^ cells/mL. 650 μL/24 well or 3ml/6 well were spun onto 1.5mm thick, uncoated glass coverslips in swing bucket rotors at 650 g for 5 minutes. Coverslips were fixed for 15 minutes with 4% formaldehyde (FA)–PBS, rinsed three times with PBS, and blocked for 1 hour in PBS containing 0.1% w/v saponin-3% bovine serum albumin (BSA, Sigma Aldrich, A7906). Samples were again rinsed three times with PBS and incubated with primary antibodies as indicated in Table S5D, diluted in PBS containing 0.1% w/v saponin-3% BSA for 1 hour at 37°C in a humidified chamber. Coverslips were rinsed three times and incubated with the appropriate species IgG (H+L) highly cross-adsorbed secondary antibody conjugated to Alexa Fluor 488 (TRAIL-R1 staining) or 568 (all others) diluted 1:1000 in PBS containing 0.1% w/v saponin-3% BSA for 1 hour at 37°C in a humidified chamber. Coverslips were washed with PBS, incubated with 4′,6-diamidino-2-phenylindole (DAPI) as directed (Invitrogen, D1306), and washed with PBS. Finally, coverslips were mounted onto glass slides using ProLong Gold antifade mountant (Invitrogen, P36934), and incubated at room temperature in the dark for 24 hours.

Confocal images were acquired on a Nikon W1 Dual Cam Spinning Disk Confocal microscope with Plan Apo λ 60x Oil objective. Widefield images were acquired in Z-stacks on a Nikon Ti-2 Widefield microscope with S Plan Fluor ELWD 40x Ph2 ADM air objective and analyzed with Imaris software for 3D colocalization of the top 10% brightest voxels per channel.

### Enzyme-linked immunosorbent assay

Anti-TRAIL ELISA was performed using a Human TRAIL/TNFSF10 Quantikine ELISA Kit (R&D Systems, DTRL00) following manufacturer-provided protocols. Briefly, 50 μL of cell supernatant or cellular lysate obtained using manufacturer provided buffers and instructions were added to each well. Cell densities at time of harvesting were 8×10^6^ to 1.2×10^7^ PEL cells, while cells were lysed at a concentration of 1×10^7^ cells/mL as per manufacturer guidelines. The included recombinant human TRAIL standard (R&D Systems, 892375) was serially diluted in 2-fold steps from 1000 to 15.6 pg/mL. Absorbance was read on a VICTOR Nivo Plate reader at 450 nm. A blank measurement taken from a well containing clean dilution buffer was subtracted from all other wells and least squares regression was performed on the measured values across all replicates to determine effective concentrations.

### In vitro TRAIL treatment

BCBL-1/Cas9 cells containing pLenti SpBsmBI Guide Hygro (AAVS1 or TRAIL-R1 #1/2) were treated with the indicated serial dilutions of recombinant human TRAIL (Sigma-Aldrich, T9701) and absolute cell numbers were measured 24 hours later by flow cytometry as described above for cumulative growth curves.

### Statistical Analysis

Statistical testing/modeling was conducted in Python using SciPy 1.6.0^88^ (simple t tests, one-way ANOVA) or Statsmodels 0.13.2 (ordinary least squares) unless otherwise indicated. Vertical bar graphs were generated using Prism 9.4.1 (GraphPad), all other figures were generated using Seaborn 0.11.2 ^89^. For groups of tests with similar hypotheses, i.e. similar comparisons in the same figure panel, false-discovery adjusted p-values were calculated using the Benjamini-Hochberg procedure implemented in Statsmodels. For two-sample t-tests, equal variances were assumed. Specific tests are indicated in the figure legends. Full code for computational and statistical analyses was made available online on Github (https://github.com/Gottwein-Lab).

## Acknowledgements

We would like to thank Dr. Joanna Shisler for sharing MCV FLIP protein expression vectors and Dr. Ethel Cesarman for sharing a control aliquot of BC-3 cells. Imaging work was performed at the Northwestern University Center for Advanced Microscopy generously supported by NCI CCSG P30 CA060553 awarded to the Robert H Lurie Comprehensive Cancer Center.

## Author Contributions

Study Design NK, SO, MM and EG. MM performed exploratory resistance screens in BC-3, which were not included in the final manuscript due to very low cFLIP dependency of this cell line. NK performed most experiments. EG performed shRNA cloning, pLX and pCW cell line construction, and shRNA resistance screens until harvest of cell pellets. SMO performed all immunofluorescence staining and analyses. ZL performed pLX and pCW vector cloning. Computational and statistical analyses were performed by NK. NK and SO put together figures, which were edited by EG. NK, SO, and EG wrote the manuscript. EG supervised all aspects of the work. All authors read and approved the final manuscript.

## Ethics Statement

This study did not require ethical approval.

## Data Availability

CRISPR screening and RNAseq data were deposited on GEO (accessions GSE210445, GSE210446). All cell lines and plasmids are available on request.

