## Supplemental Figures and Legends for "CRISPR Screens Identify Novel Regulators of cFLIP Dependency and Ligand-Independent, TRAIL-R1-Mediated Cell Death"

### Supplemental Material

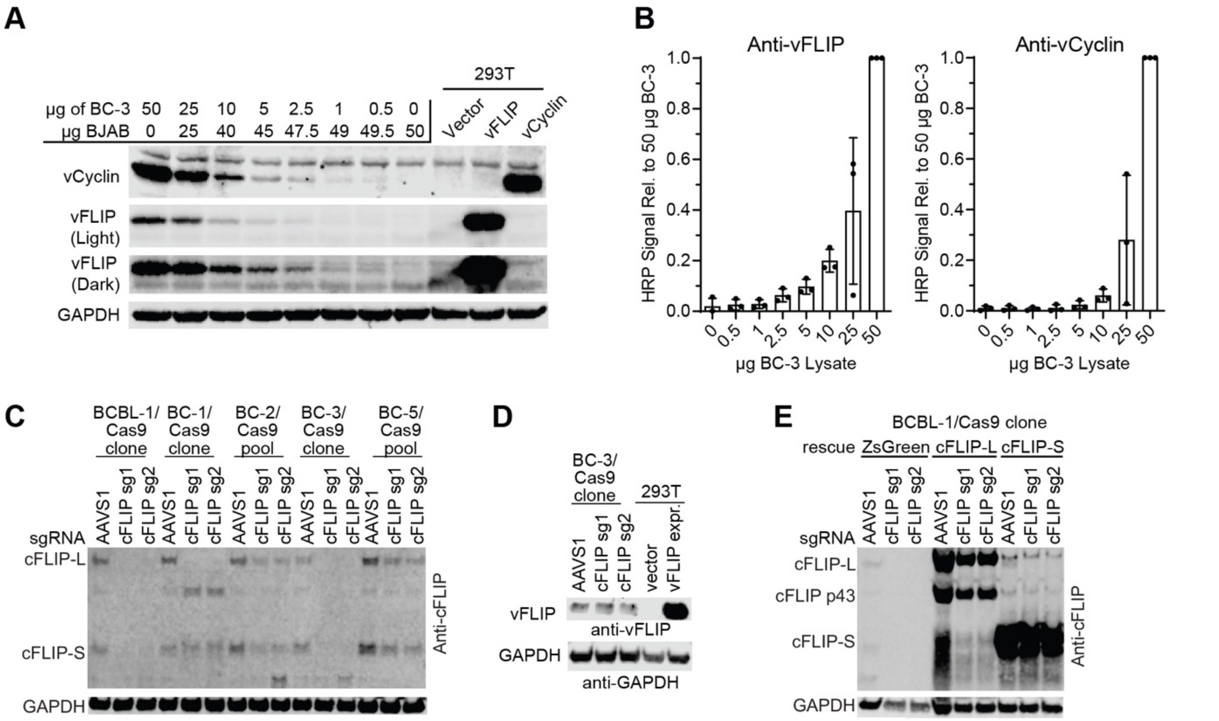


**Supplemental Fig. 1** Sensitivity of vFLIP/vCyc Westerns and cFLIP KO controls for Fig. 1 **A.** BC-3 and BJAB lysates were harvested, and the indicated amounts of proteins were mixed to achieve a total of 50 μg per lane. Lysates from 293T cells transfected with vCyc or vFLIP expression vectors were included as additional controls (n=3). **B.** Band intensity was quantified for independently harvested lysates, as in panel (A), via densitometry. Error bars indicate SD (n=3). **C.** Lysates were harvested on day 3 after transduction with the indicated sgRNAs during the cumulative growth curves (shown in Fig. 1B) and cFLIP Western blotting was performed to confirm knockout efficiency (n=3). **D.** The lysates from BC-3 in panel C were also probed for vFLIP expression via Western blots (n=2). **E.** As in panel (C) but accompanying Fig. 1C (n=3).

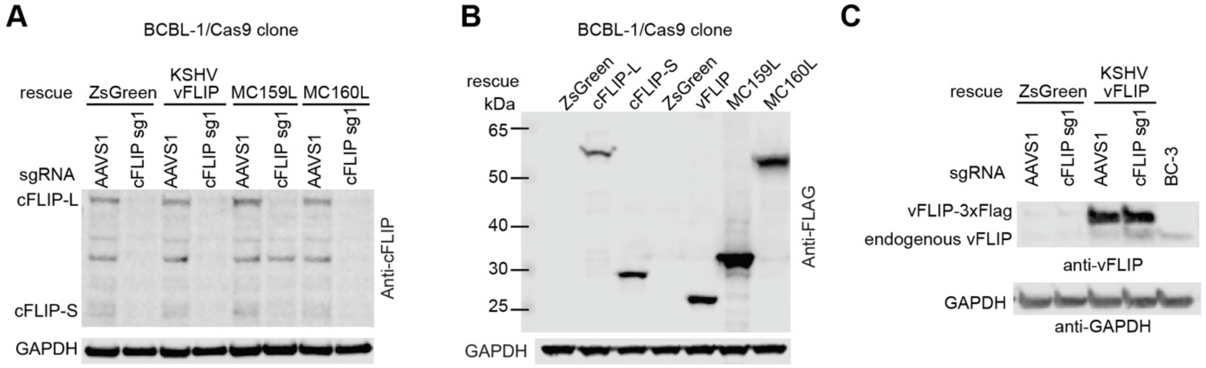


**Supplemental Fig. 2** Expression controls for Fig. 2. **A.** Lysates collected 2 days after transduction during the cumulative growth curves shown in Fig. 2B were subjected to Western blot analysis of cFLIP expression (n=3). **B.** Lysates were harvested from the ectopic expression lines utilized for experiments in Fig. 1C and 2B and subjected to Western blot analysis of total FLAG epitope levels to facilitate comparison of overall expression levels of the various FLIP proteins (n=1). **C.** ZsGreen control and vFLIP ectopic expression lysates from panel (A) were additionally probed by Western blot for vFLIP expression (n=2).

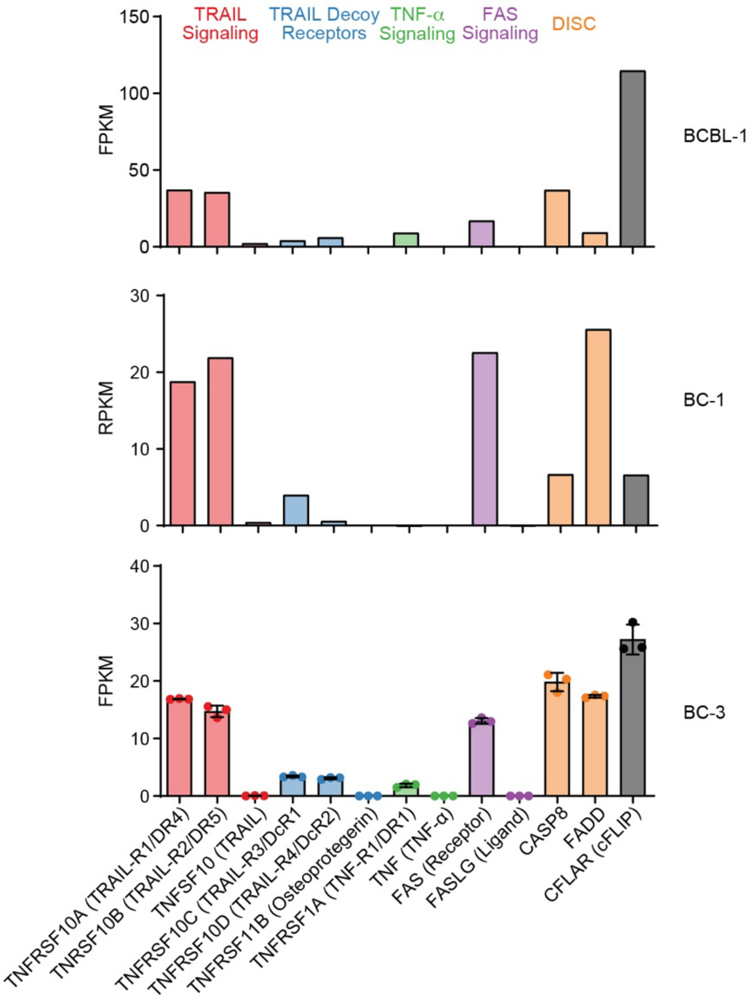


**Supplemental Fig. 3**. Quantification of mRNA levels for selected death signaling genes. Reads or fragments per kilobase per million were calculated for the three PEL cell lines indicated. Data for BC-3 and BCBL-1 were obtained from datasets previously published by us and others[64, 90], while our BC-1 data represent new data. RPKM is shown for single-ended sequencing data, while FPKM is shown for paired-end data. For BC-3, error bars indicate SD (n=3, samples harvested side-by-side). We note that BCBL-1 represented a DMSO treated control.

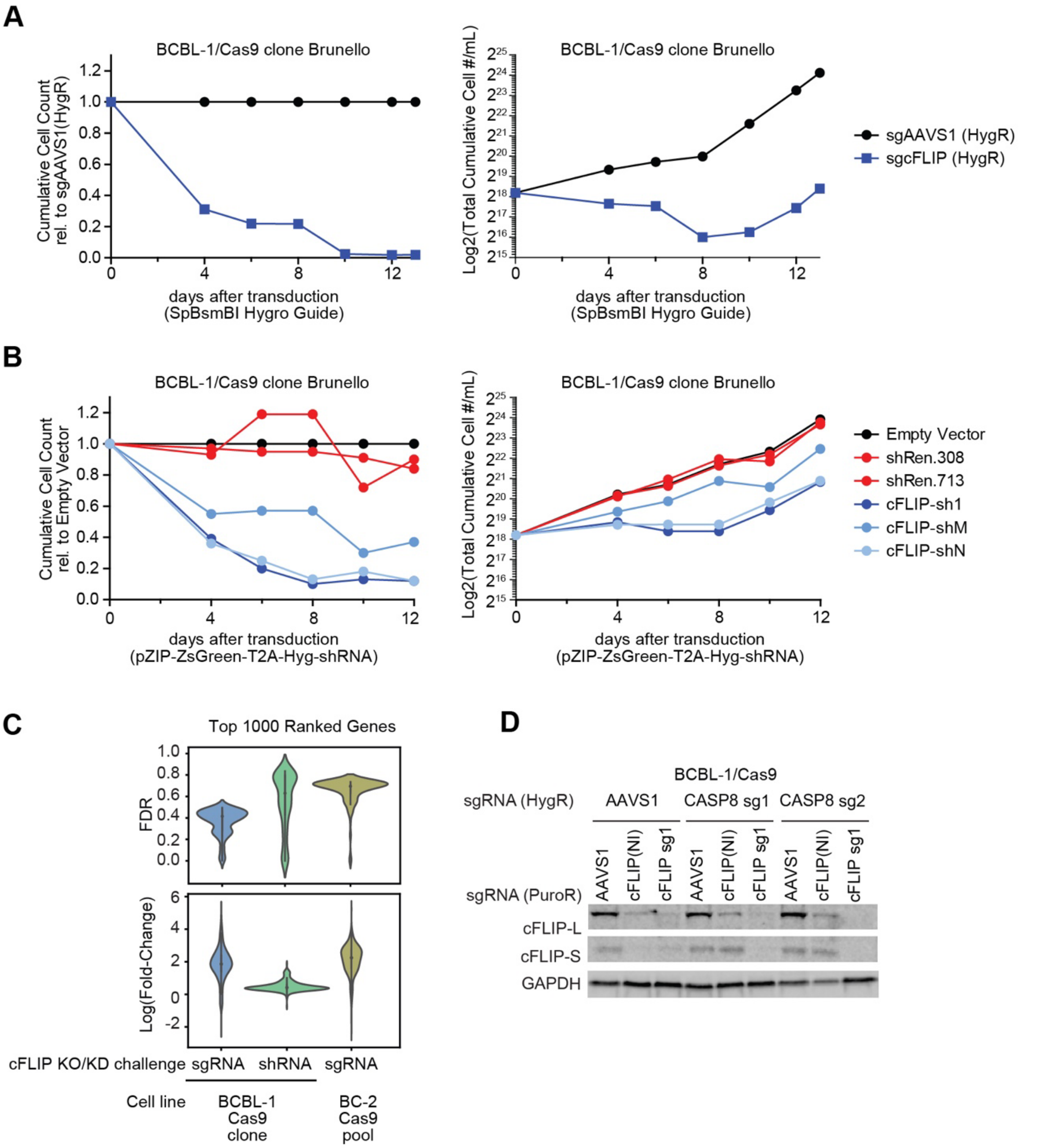


**Supplemental Fig. 4**. Additional data supporting cFLIP resistance screens and CASP8 knockout experiments. BCBL-1/Cas9 cells were transduced with the lentiviral sgRNA library Brunello, followed by the indicated **A.** sgRNA or **B.** shRNA constructs. Cumulative cell counts relative to the appropriate control (left) or log2-transfomed raw cumulative counts (right) are shown starting on the day of the secondary transduction (sgRNA/shRNA challenge). Error bars indicate SD (n=3). **C.** The top 1000 ranked genes for each screen were selected by MAGeCK/RRA-rank. The FDR and fold-change values for these genes were then plotted with inter-quartile range (solid line) and Gaussian kernel density estimates (colored contours). **D.** Lysates harvested 3 days after transduction with the indicated sgRNA during the cumulative growth curves displayed in Fig 4B. Lanes marked cFLIP (NI, not included) represent a third, cFLIP sgRNA not included elsewhere in our study, since this guide resulted in only inefficient editing.

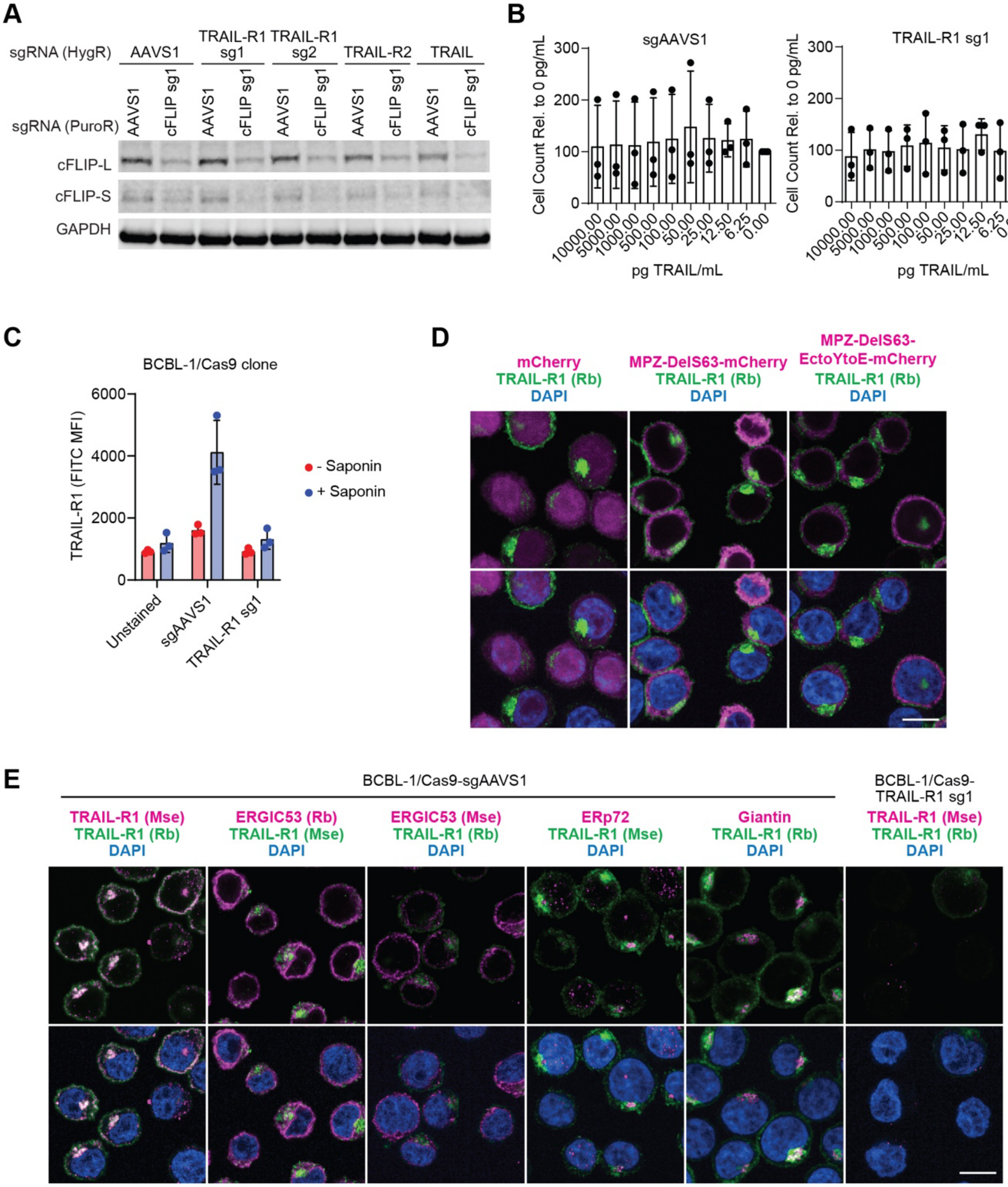


**Supplemental Fig. 5**. Additional data supporting TRAIL-independent TRAIL-R1 signaling in Fig. 5. **A.** Lysates were collected 2 days after sgRNA transduction during the cumulative growth curves shown in Fig. 5B and cFLIP knockout efficiency was established by Western blot. **B.** BCBL-1/Cas9 cells transduced with control (AAVS1) or TRAIL-R1 (sg1) sgRNAs were treated with the indicated concentrations of TRAIL ligand and total cell counts were quantified 24 hours later. Error bars indicate SD (n=3 independent repeats). There were no significant differences between doses for both cell lines, tested by one-way ANOVA (p=.99 for both). **C.** Flow cytometry on TRAIL-R1 stained cells was performed as in Fig. 5D. Displayed is the mean fluorescence intensity (MFI) across 3 independent repeats, error bars indicate SD. **D.** Representative images of TRAIL-R1 colocalization with mCherry and mCherry-tagged MPZ mutants are shown. TRAIL-R1 is shown in green with mCherry in magenta to depict overlapping signal as white. Scale bar = 10μm. **E.** Representative images of TRAIL-R1 and all compartment markers are shown. TRAIL-R1 is shown in green with all other markers in magenta to depict overlapping signal as white. Scale bar = 10μm.

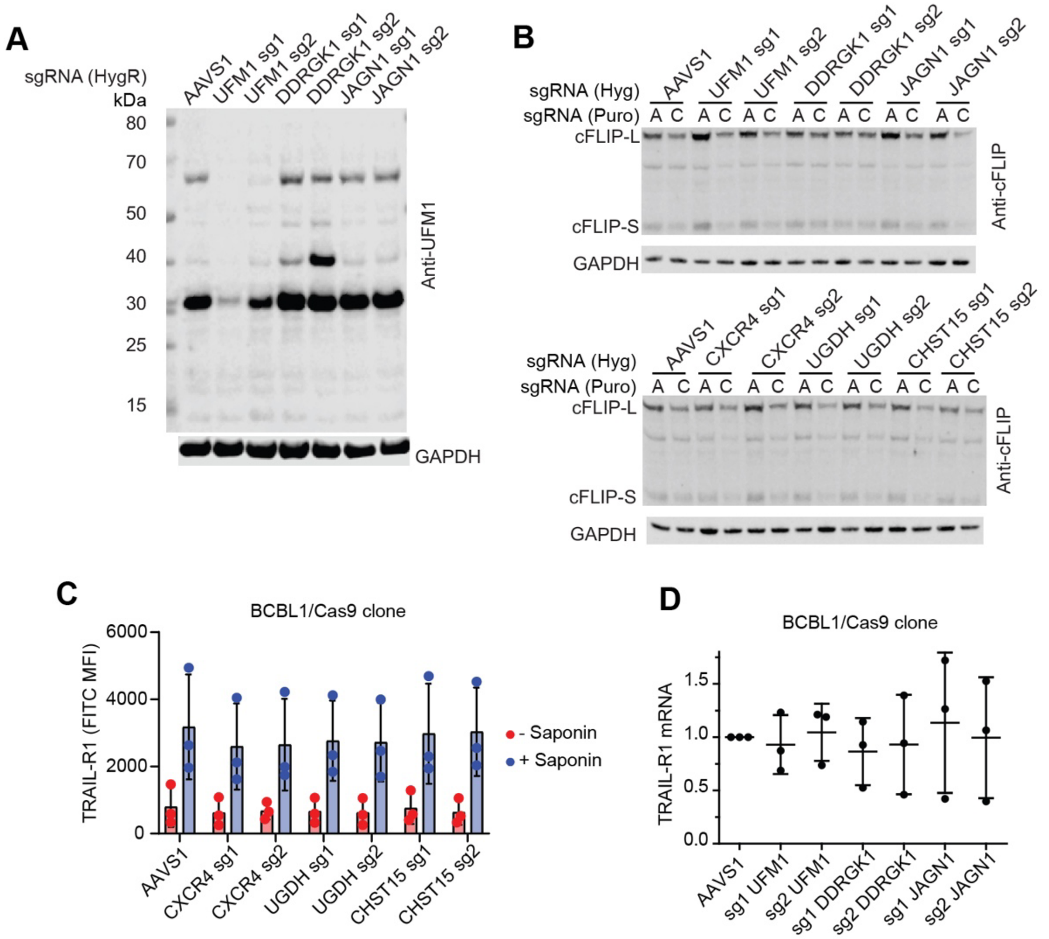


**Supplemental Fig. 6**. Additional data supporting UFMylation, chondroitin sulfate synthesis, JAGN1, and CXCR4 knockout experiments. **A.** Lysates from a subset of single-gene knockout pool cell lines from Fig. 6A were assessed for UFM1-conjugates by Western blot analysis. Unconjugated UFM1 is not shown as its low molecular weight (12 kDa) leads to co-migration with the loading dye front. **B.** Lysates were collected 2 days after transduction during the cumulative growth curves shown in Fig. 6A and cFLIP knockout efficiency was confirmed by Western blot. Incomplete cFLIP KO is due to performing this and similar Westerns before excessive cell death is observed. **C.** The indicated subset of single gene knockout cell lines from Fig. 6 were stained for TRAIL-R1, and flow cytometry was performed in parallel to the experiments shown in Figs. 5D and S5C. Displayed is the MFI across 3 independent repeats, error bars indicate SD. The lack of significant differences between cell lines within permeabilization groups was tested by one-way ANOVA ($p=0.99$ for both). **D.** Real-time PCR was performed on cDNA isolated from the indicated single-gene knockout pool cell lines. Expression is quantified relative to an endogenous control (B2M) and the sgAAVS1 control. Each point represents the average of 3 technical replicates on a single plate ($n = 3$ independent RNA preparations). Lack of significant differences between cell lines was tested by one-way ANOVA ($p=0.98$).

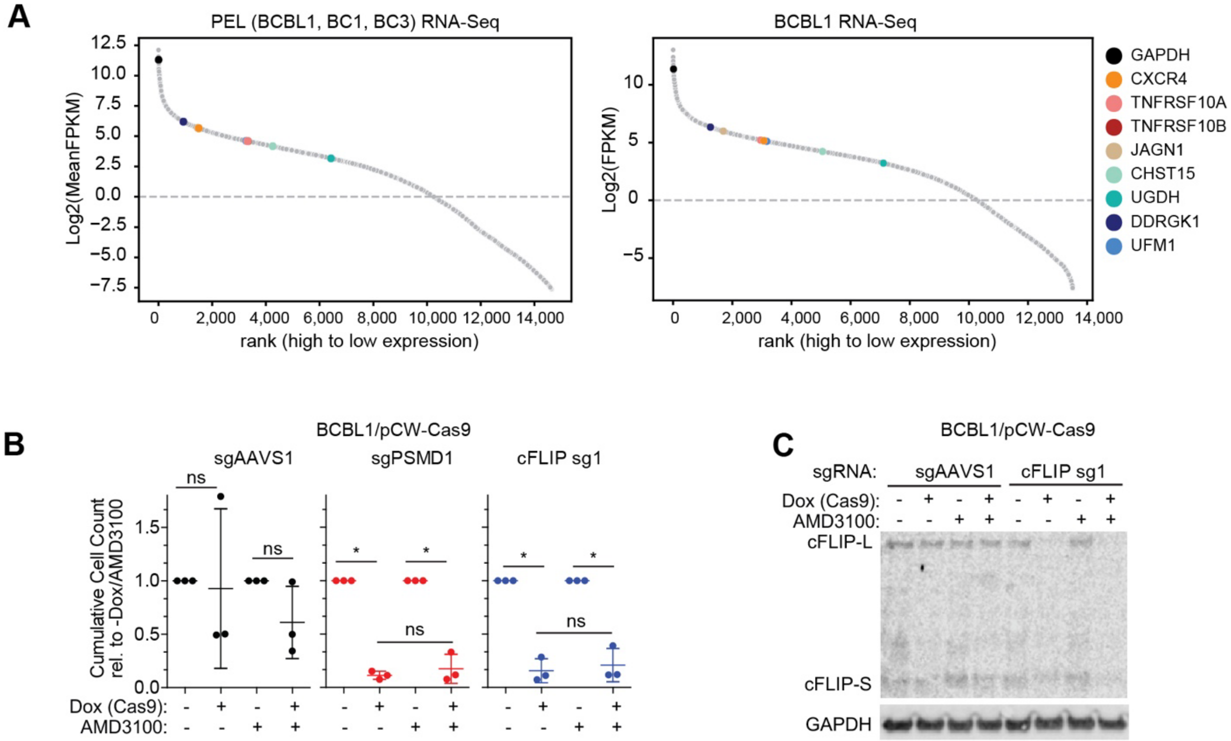


**Supplemental Fig 7.** Expression of CXCR4 and chemical inhibition of CXCR4 signaling. **A.** CXCR4 and other genes chosen for validation in Fig. 6 are well expressed in the RNA-seq datasets from Fig. S3, while CXCL12 expression is not detected (not shown). **B.** BCBL-1 cells with Dox-inducible Cas9 expression (BCBL-1/pCW-Cas9) were transduced with the indicated sgRNAs, selected, and treated with the indicated combinations of Dox and 25µM AMD3100. Cumulative cellular growth curves over seven days were performed as described above (n=3 independent repeats). Lack of significant differences was tested by one-tailed, two-sample t-tests, FDR-adjusted p-values are listed in Table S6. **C.** Lysates were collected 5 days after the start of treatment in E) and cFLIP knockout efficiency was confirmed by Western blot (n=2).

**Supplementary Data Legends**

**Supplementary Data 1**. Gene-level output from MAGeCK-RRA cFLIP depletion resistance screens. The gene-level results of the three resistance screens described in Fig. 3 and S4 are provided as separate Excel tabs. Results represent status in cFLIP depleted cells relative to their appropriate controls, such that enrichment values represent genes with guides that are more abundant following cFLIP depletion.

**Supplementary Data 2**. Summary of high-confidence enrichment targets from cFLIP depletion resistance screens. Data for the intersection of the top 150 ranked genes in both BCBL-1 screens (plus UFL1, which narrowly misses this cutoff in our BCBL-1 sg-cFLIP resistance screen) are shown. The following data are shown for each gene: 1) gene name and a short functional description; 2) statistical enrichment values output by MAGeCK-RRA for each cFLIP depletion resistance screen, as found in Supplementary Data 1; 3) median negative FDR of depletion across 8 PEL CRISPR screens, as published previously[7]; 4) data collected from the Q2 2022 release of the Cancer Dependency Map regarding the percentage of cell lines with a significant dependency on the indicated gene and whether than gene was classified as common essential or strongly selective.

**Supplementary Data 3.** DAVID pathway analysis of high-confidence enrichment targets from cFLIP depletion resistance screens. Data for the intersection of the top 150 ranked genes in both BCBL-1 screens (23 in total—UFL1 not included in this case) were input into the DAVID pathway analysis tool, utilizing all guides detected in either BCBL-1 screen as a background list. Full output for the three broadest categories of gene ontology terms (biological processes, cellular components, and molecular functions) is provided, including which of our enriched genes occur within the indicated pathways. Pathways selected for inclusion into Fig. 3C are bolded and represent the top non-redundant pathway in each case.

**Supplementary Data 4.** NGS-based validation of CRISPR editing of selected knockouts. The first tab represents a summary describing each single guide knockout pool on a different row. Columns include: 1) summary statistics regarding assembled contigs used for custom variant analysis (total read pairs, total variants/unique contigs, and the percentage of total reads mapping to assembled contigs); 2) rates of different variant classifications (indel, frameshift, synonymous, missense, and nonsense) in custom analysis pipeline as a percentage of all contig-mapped reads; 3) the expected and predicted length of peptides based on stop codon position, provided as three measures of central tendency across all reads mapped to contigs (mean, median, and mode); 4) rates of modified reads and frameshifted reads as predicted by CRISPResso2 (note that CRISPResso2 provides frameshift rates relative to modified reads, not total reads) [86]. Complete results for each contig are provided in a separate Excel tab for each knockout pool.

**Supplementary Data 5.** List of reagents detailed in Material & Methods. The following types of reagents are listed in individual tabs: **A.** PCR primers used for cloning and modification of original vectors **B.** Gene fragments/blocks used for cloning original vectors **C.** List of single guide RNAs and the corresponding oligonucleotides used for cloning into the indicated lentiviral guide vectors **D.** Antibodies used for Western blots, flow cytometric analysis, or immunofluorescence. **E.** Library preparation primers used for sequencing on the Illumina NextSeq platform along with barcode sequences, while those used for the HiSeq platform were previously described[7] **F.** Primers used for amplicon-based CRISPR variant sequencing.

**Supplementary Data 6.** Statistical testing output and summary for cumulative growth curve experiments. Each Excel tab corresponds to the indicated figure/panel. Description of full statistical testing methodology is provided in Materials and Methods.
